# Precise metabolic dependencies of cancer through deep learning and validations

**DOI:** 10.1101/2025.02.26.640327

**Authors:** Tao Wu, Xiangyu Zhao, Yu Zhang, Die Qiu, Kaixuan Diao, Dongliang Xu, Weiliang Wang, Xiaopeng Xiong, Xinxiang Li, Xue-Song Liu

**Author notes:** These authors contributed equally to this work (Tao Wu, Xiangyu Zhao, Yu Zhang).

## Abstract

Cancer cells exhibit metabolic reprogramming to sustain proliferation, creating metabolic vulnerabilities absent in normal cells. While prior studies identified specific metabolic dependencies, systematic insights remain limited. Here we build a graph deep learning based metabolic vulnerability prediction model “DeepMeta”, which can accurately predict the dependent metabolic genes for cancer samples based on transcriptome and metabolic network information. The performance of DeepMeta has been extensively validated with independent datasets. The metabolic vulnerability of “undruggable” cancer driving alterations have been systematically explored using the cancer genome atlas (TCGA) dataset. Notably, *CTNNB1* T41A activating mutations showed experimentally confirmed vulnerability to purine/pyrimidine metabolism inhibition. TCGA patients with the predicted pyrimidine metabolism dependency show a dramatically improved clinical response to chemotherapeutic drugs that block this pyrimidine metabolism pathway. This study systematically uncovers the metabolic dependency of cancer cells, and provides metabolic targets for cancers driven by genetic alterations that are originally undruggable on their own.

## Introduction

Metabolic reprogramming of tumor cells is closely related to several processes associated with tumor progression, including the initiation of transformation, proliferation and metastasis of tumor cells, and has been considered one of the hallmarks of cancer^1^. Metabolic reprogramming improves the adaptability of tumor cells and gives them a selective advantage over other cells, increasing the probability of tumor cells surviving under stress conditions. In turn, these metabolic adaptations make cancer cells dependent on specific metabolic pathways, thus creating vulnerabilities that can be therapeutically explored.

Genetic alterations are major driving forces for cancer, and some driving genetic alterations, such as *EGFR*-L858R, *BRAF*-V600E, have created drug targets for treating cancers that are dependent on these drivers. However there are many recurrent cancer driving events, such as genetic alterations in *CTNNB1, MYC, TP53*, that generate mutated proteins lacking accessible hydrophobic pockets, in which small molecules can bind with high affinity and thus are usually termed “undruggable”^2^ or “difficult to drug” directly. Meanwhile, many cancer driving genetic events can lead to metabolic alterations. The following are examples: *MYC* overexpressed gastrointestinal cancer cells show specific vulnerability to thymidylate synthase (TYMS) inhibition^3^. *KRAS/LKB1* co-mutant lung cancer cells show metabolic vulnerability to hexosamine biosynthesis pathway (HBP) inhibition^4, 5^. *TP53* mutant triple negative breast cancer has been reported to show specific sensitivity to GPX4 inhibition or knockdown compared with *TP53* wild type cells^6^. Thus, targeting these cancer-driving genetic alterations indirectly by exploiting metabolic dependency could be an appealing strategy. However, systematic investigations of these metabolic vulnerabilities for cancer cells are still lacking.

The exploration of tumor cell-specific metabolic dependencies by *in vitro* or *in vivo* studies is time-consuming and laborious. Cellular metabolism forms a natural graph or network architecture, with different metabolic enzymes connected through metabolites. The metabolic vulnerability of each specific sample is dependent on the sample status and the metabolic pathway context. Sample states can be modeled by gene expression profiles, while relationships among complex metabolic pathways can be modeled by metabolic networks. Graph Neural Networks (GNNs) are a powerful approach to extract information from graph-structured data, such as biological networks, including protein-protein interaction network ^7^ and gene regulation network^8^. Besides, recent developments of large scale CRISPR-based screening data, coupled with data of cancer cell lines subjected to comprehensive omics characterization, have provided critical resources for identification of genes essential for cancer cell survival^9^. Together, advances in graph deep learning and accumulation of data make it possible to model metabolic dependencies and perform *in silico* screening to accelerate the discovery and validation of cancer-specific metabolism dependency targets.

Here we build a deep learning based algorithm “DeepMeta” that can accurately predict the metabolic vulnerability of individual cancer patient based on the transcriptome and metabolic enzyme network information. We applied this model to the cancer genome atlas (TCGA) tumor samples and found that nucleotide metabolism and glutathione metabolism have universal metabolic dependency. Applying to TCGA samples, the model predicted the metabolic dependence for common undruggable cancer drivers, including genetic alterations in *TP53*, *CTNNB1*, and *MYC*. We performed experimental validations for the model-predicted metabolic dependencies of important cancer genes and found that *CTNNB1* activating mutation leads to a dependency on pyrimidine and purine metabolism, and chemically or genetically inhibit these metabolic pathway can specifically block the proliferation of cells with *CTNNB1* activating mutation.

## Results

### The DeepMeta framework

Here, we present the DeepMeta framework to predict the metabolic gene dependence based on the metabolic pathway information characterized by enzyme networks and sample status information defined by gene expression profile, shown as two inputs in **Figure 1A**. We used the graph attention network^10^ (GAT) module to extract information from the sample-specific enzyme network to obtain the embedding of the metabolic enzyme gene node, and used the fully connected neuron network module to extract information from the expression profiles of samples to get the sample state embedding (see **STAR Methods**). Here, the features of metabolic gene node in metabolic enzyme network was implemented by binary vector which denotes the involvement of the gene in chemical and genetic perturbation (CPG) gene sets^11^. These two parts of information are then merged to predict metabolic dependencies for each specific sample. DeepMeta was trained on cell line CRISPR-Cas9 screening data of DepMap^12^ (23Q2 version).

**Figure 1:**
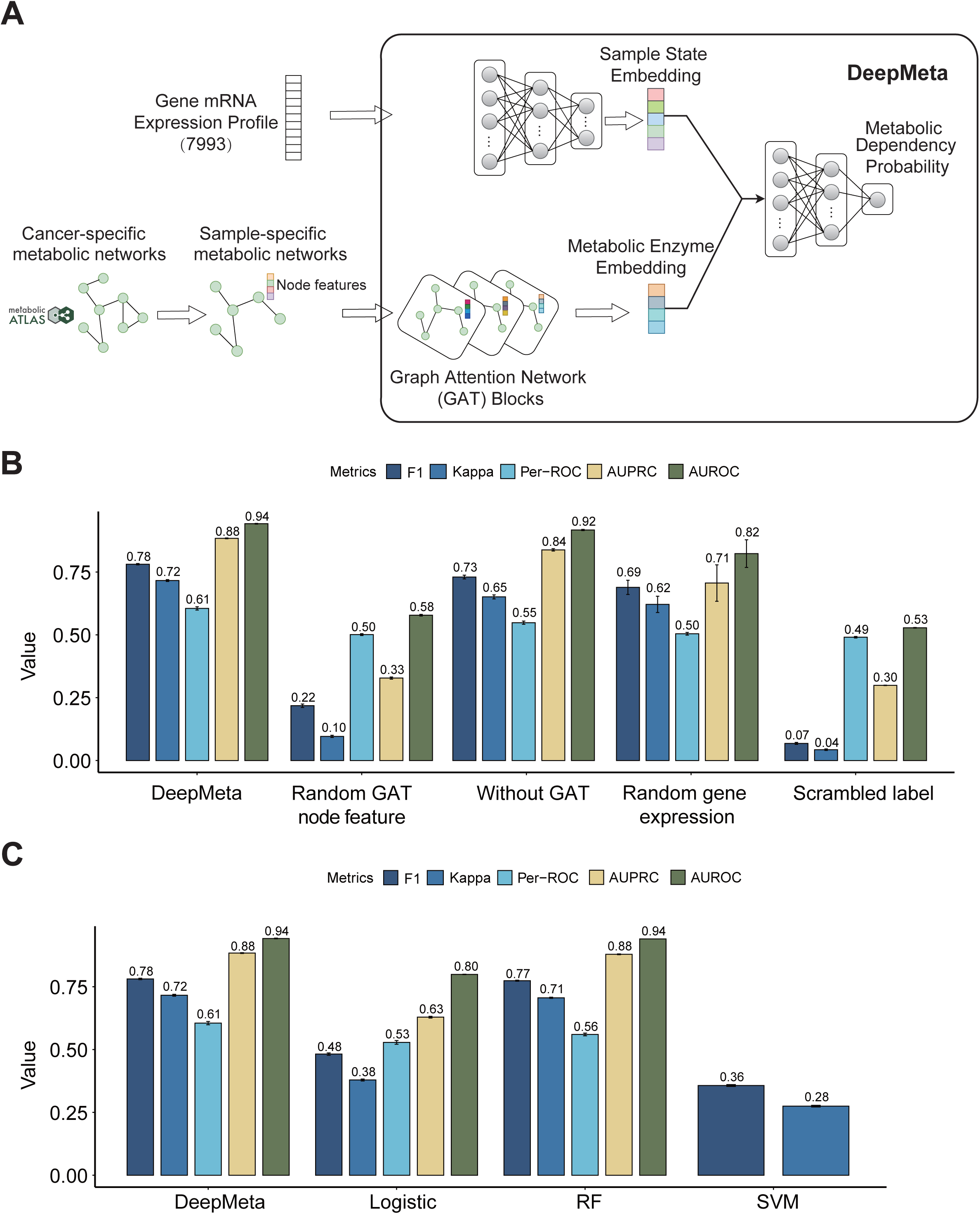
Framework of DeepMeta and model comparison. **A,** Illustration of DeepMeta framework. DeepMeta has two modules to handle different kinds of input. The GAT module is used to extract information from the sample-specific enzyme network (SEN) to obtain the embedding of the metabolic enzyme gene node, and the fully connected layer module is used to extract information from the expression profile of samples to get the sample state embedding. SEN was constructed by removing genes in the cancer-specific enzyme network that are not expressed in the sample. These two parts of information are then merged to predict the metabolic dependencies for each sample. **B,** Ablation experiments to access the impact of different parts of DeepMeta model on the performance. Performance comparison of different models using random gene node features, without the enzyme network, random gene expression, or dataset with scrambling label as input. All models were trained in ten-fold cross-validation manner and metrics in every test fold were reported. **C,** Model performance comparison of DeepMeta with the three classic machine learning models. The metrics include F1 value, Kappa value, the area under the receiver operating characteristic curve (AUROC) and the area under the precision-recall curve (AUPRC). We also evaluated the prediction performance of individual genes by AUROC (termed per-ROC). The number on top of bar indicates mean value of the metric in ten test folds.

To access the impact of different parts on the performance of the model, we conducted ablation experiments, and train new models using random metabolic node features, without the enzyme network or random sample expression as input, then compared the performance of these model with DeepMeta using ten-folds cross validation (see **STAR Methods**). Compared with models using random metabolic node features and random gene expression, DeepMeta shows a marked performance improvement (**Figure 1B**). We found that performance of the model using random metabolic node features decreased more severely than the model using random gene expression, suggesting that CPG node features are more important for predicting metabolic dependencies than the gene expression of samples. DeepMeta also has a performance improvement over models that do not use the metabolic enzyme network as input (i.e. without the GAT module), indicating that the information extracted by the GAT is beneficial for the model to learn metabolic dependencies. We also performed scrambling label experiment as negative control to verify that the achieved performance was not by chance. Besides, our model achieved an average ten-fold per-ROC (AUROC of individual gene) of 0.61, much higher than expected by random chance using the scrambled-label method (average per-ROC of 0.49, **Figure 1B**). Among these metabolic genes, the proportion of genes with an ROC exceeding 0.5 is about 85% (**Figure S1**), indicating that our model has good predictive performance for most metabolic genes.

We compared the model performance with three machine learning (ML) models (see **STAR Methods**). Compared with logistic regression and support vector machine (SVM) models, DeepMeta showed a marked performance improvement (**Figure 1C**). Although our model showed small improvement compared with random forest (RF) in the ten-fold cross validation analysis, the generalization ability of RF is much lower compared with DeepMeta (**Figure S2**, the description of independent datasets is available in the next section). Furthermore the dimensionality reduction preprocessing operation also makes the RF model difficult to interpret^13^. Overall, our DeepMeta model exhibits the better performance when compared with classical machine learning models.

### Model performance evaluation in independent datasets

In the test dataset (76 cell lines and 1010 genes), DeepMeta achieved the ROC of 0.94 (**Figure 2A**) and other metrics also reached relatively high performance **(Figure S3**), illustrating our model enables accurate prediction of metabolic dependencies. We further validated DeepMeta performance using five independent datasets: 1) Another CRISPR screen data conducted by the Sanger institute using a different CRISPR library^14^, containing 200 cell lines overlapped with training data and 25 unique cell lines; 2) RNAi screen data, which apply DEMETER2^15^ to the combination of three large-scale RNAi screening datasets: the Broad institute project Achilles^12^, Novartis project DRIVE^16^, and the Marcotte et al^17^ breast cell line dataset. This dataset contain 417 overlapped cell lines and 160 unique cell lines; 3) The Childhood Cancer Model Atlas^18^ (CCMA) dataset which performs CRISPR-Cas9 screen in 110 paediatric solid tumour cell lines and reports Z scores; 4) Pemovska et.al^19^ drug screen dataset, which performs metabolic drug library (contains 243 compounds) screen in 15 cancer cell lines (14 cell lines with gene expression data) and reports AUC (area under the curve) scores, larger AUC score indicates a stronger killing effect on the cell; 5) PRISM repurposing dataset, which contains the results of pooled-cell line chemical-perturbation viability screens for 4,518 compounds screened against 578 cell lines and the drug effect score is represented by log fold change (ratio of cell viability between treatment and control). DeepMeta achieved high AUROC scores in Sanger and RNAi dataset, both for overlapped and unique cell lines (**Figure 2B-C**). The slightly low performance metrics in RNAi dataset (**Figure S3**) could be partly due to the low correlation between the RNAi and the CRISPR dataset (**Figure S4**). Although the model only achieved a moderate AUROC score in the CCMA dataset, there is a significant difference in the Z scores between different predicted classes (**Figure 2D**). For each cell line-gene pair in Pemovska dataset, we first obtain the maximum AUC value of all drugs targeting a specific metabolic gene, then rank the genes in each cell line based on this AUC value and assign ranks. We showed the overlap of the top 5 genes predicted by DeepMeta and the genes with top 3 AUC rank in each cell line (**Figure 2E**). Interestingly, we found that 11 of the 14 cell lines contain overlap genes and the AUC rank of the most overlapped genes is 1 (8/11, median AUC value of these overlapped gene is 0.85). This result indicates that DeepMeta predictions can be recapitulated by actual drug screening experiments. In PRISM dataset, we found that log fold change values of drugs targeting gene with predicted positive label were significantly lower than drugs targeting gene with predicted negative label (**Figure S5**), indicating drugs targeting DeepMeta predicted positive genes have stronger tumor cell killing effects. In summary, DeepMeta shows good performance and generalization ability on different independent datasets.

**Figure 2:**
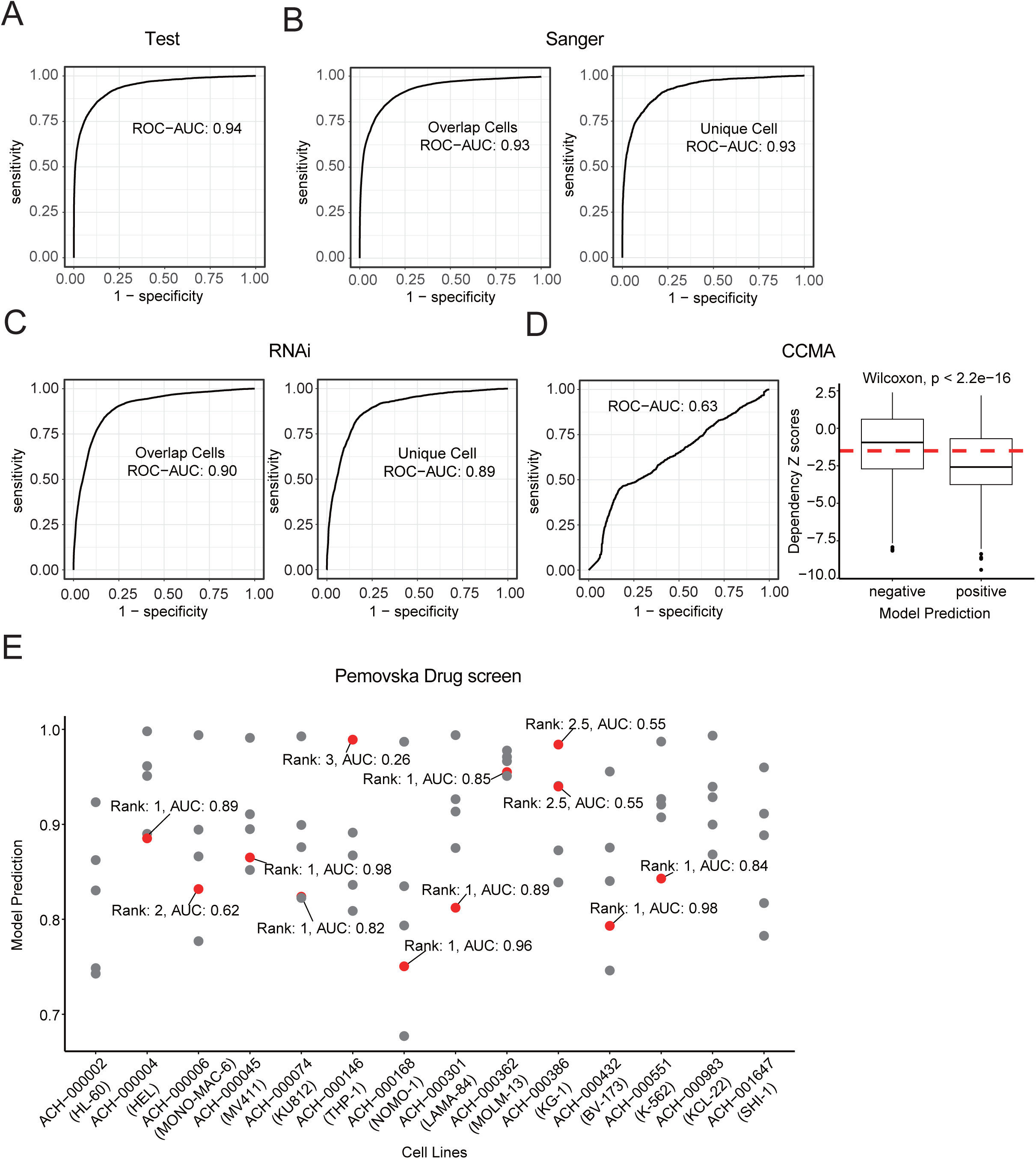
DeepMeta performance validation using additional datasets. **A-D,** DeepMeta performance shown as ROC curves and AUROC values on the test set (**A**), and independent validation datasets: Sanger dataset (**B**), RNAi dataset (**C**), and CCMA dataset (**D**). In CCMA dataset, Z score was used to quantify gene dependency. Thus we also compare Z scores between different predicted categories. **E,** Performance validation in Pemovska drug screen dataset. In each cell line, the metabolic enzyme gene was assigned a rank value based on AUC score of drugs targeting this metabolic enzyme. The top 5 metabolic genes predicted by DeepMeta are shown and the top 3 experimentally validated metabolic genes were labeled in red color.

Explaining the complex deep learning model helps us to understand its learning capabilities. The GAT model, due to its utilization of attention mechanisms during model training, is a naturally interpretable model. Thus, we extracted attention weights from the GAT model to determine the importance of a metabolic enzyme’s neighbor enzymes for its prediction (**Figure S6A**). For each pair of metabolic enzyme genes, we computed the correlation of their post-Chronos dependency scores^27^ and compared the difference in neighbor importance scores between different correlation groups. In comparison to groups with low dependency correlation (R < 0.2), those with higher dependency correlation (R > 0.8) exhibited significantly elevated neighbor importance scores (**Figure S6B**). This demonstrates that our model could utilize attention mechanism to learn the metabolic dependency associations between genes. We show the metabolic enzyme genes that are crucial for predicting the metabolic dependency of the *GPX4* in **Figure S6C** as an example. The model reveals that the most essential gene for predicting *GPX4*’s metabolic dependency is *GSR*, which aligns with existing study^20^. The distribution of important genes for predicting the metabolic dependency of some additional metabolic genes can be found in **Figure S7.**

With the availability of genome-wide loss-of-function screening datasets, predicting the cancer dependency become a reality. DeepDep^21^ is the transfer learning based model for predicting cancer dependency. However, DeepDep is not designed for metabolic dependency prediction, and the complex metabolic enzyme network information has not been considered in this model. We evaluated the performance of DeepDep model using independent datasets, including the Pemovska *et al* drug screening dataset, as well as the CCMA dataset. The AUC of the overlapping genes between the top 5 genes predicted by DeepDep and genes with top 3 AUC rank in each cell line is significantly lower than that of DeepMeta model in the drug screening dataset (the median AUC of the DeepDep model is 0.57, and the median AUC of the DeepMeta model is 0.85, **Figure S8A**). In the CCMA data set, the AUROC of the DeepDep model is only 0.51 (**Figure S8B**); there is no significant difference of Z values between the positive and negative samples predicted by DeepDep (**Figure S8C**). The median correlation between the actual Z value and predicted scores is only 0.4 (**Figure S8D**). These data demonstrate that the generalization ability of our model significantly outperforms DeepDep.

### Pan-cancer metabolic dependency

We applied the DeepMeta to TCGA samples, including 8937 samples of 25 cancer types, to predict metabolic dependencies at the pan-cancer level. The correlation of the gene prediction values between the cell line and tumor tissue was highly consistent (the median correlation was 0.6, **Figure S9**). For each KEGG metabolic pathway, we showed the distribution of the predicted dependency probability in TCGA samples (**Figure S10**). Then we performed enrichment analysis of metabolic pathways dependency (see **STAR Methods**). Nucleotide metabolism (including purine metabolism and pyrimidine metabolism) and glutathione metabolism showed the top significant dependency difference between cancer and normal, both in pan-cancer (**Figure 3A**) and individual cancer types (**Figure S11**). To mitigate the potential impact of pathway gene set size on enrichment results, we conducted permutation test (see **STAR Methods**). The distribution of percentage differences of samples showing dependency to metabolic pathways between TCGA cancer and normal tissues confirmed that nucleotide metabolism and glutathione metabolism are the universal cancer specifically dependent metabolic pathways (**Figure S12**). Furthermore, the metabolic dependency difference of these pathways is still significant when controlled the percentage information of stromal cells calculated by the CIBERSORT (**Figure S13**).

**Figure 3:**
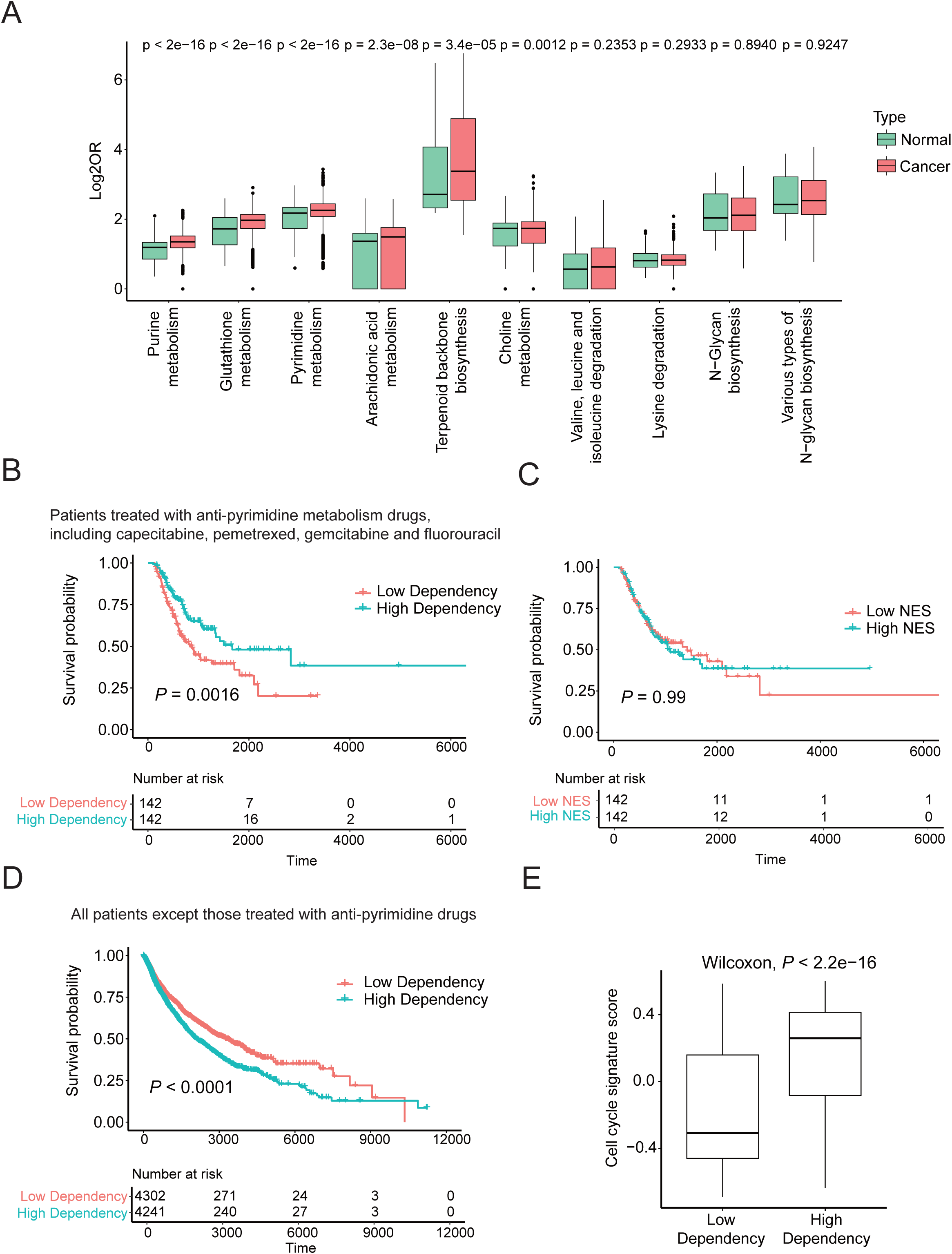
Pan-cancer metabolic dependencies and clinical validations. **A,** Compare the dependent metabolic pathway in cancer vs normal samples in TCGA pan-cancer dataset. Showing is the comparison of odds ratio (OR) values (obtained by one tail Fisher test) of metabolic pathways between TCGA cancer and corresponding normal samples, and the metabolic pathways are ranked from left to right according to *P* values. **B,** Kaplan–Meier overall survival curves show the comparison between different cancer patients stratified by the dependency in pyrimidine metabolism pathway. These patients were treated with four anti-pyrimidine drugs, including capecitabine, pemetrexed, gemcitabine and fluorouracil. The cutoff between “high dependency” and “low dependency” patient group is the median OR value for pyrimidine metabolism pathway dependency. **C,** Kaplan–Meier overall survival curves show the comparison between different cancer patients stratified by enrichment of pyrimidine metabolism pathway. These patients were same as (B). We used gene set enrichment analysis (GSVA) to calculate normalized enrichment score (NES) of pyrimidine metabolism pathway for each sample based on gene expression data. Then samples were stratified by median value of NES to NES low and high group. **D,** Kaplan–Meier overall survival curves show the comparison between different cancer patients stratified by the dependency in pyrimidine metabolism pathway. These patients are all TCGA patients except patients treated with anti-pyrimidine drugs. In this survial analysis all cancer types were pooled together, individual cancer type analysis and Cox analysis considering cancer type, age and gender as covariate were shown in Figure S13. **E,** Compare the cell cycle signature score difference between the two group of patients with different pyrimidine metabolism dependency. The cell cycle signature score is calculated by gene set variation analysis (GSVA) method.

To investigate the potential clinical applications of DeepMeta, we divided patients from the TCGA dataset which received anti-pyrimidine metabolism drug treatment (including capecitabine, pemetrexed, gemcitabine and fluorouracil) into two groups based on the dependency of pyrimidine metabolism pathway and calculated the survival difference between the two patient groups. We found that among patients who received anti-pyrimidine metabolism drug treatment, those predicted by DeepMeta to have a higher dependence on pyrimidine metabolism had dramatically improved survival (**Figure 3B, Figure S14A**), and this survival difference is still significant after considering cancer type, age and gender covariates. (**Figure S14B**). While there was no survival difference when the samples were stratified based on the expression enrichment of pyrimidine metabolic pathway (**Figure 3C**). Interestingly, among patients who had not been treated by anti-pyrimidine drugs, the group with higher dependency exhibits significantly worse prognosis (**Figure 3D** and **Figure S14B**). In these patients, we compared the enrichment of cell cycle-related gene signature (through gene set variation analysis). The results showed that patients with higher dependency on pyrimidine metabolism pathway had more active expression of cell cycle-related genes (**Figure 3E**), indicating the tumors which are more dependent on pyrimidine metabolism may have stronger proliferation phenotype, thus worse prognosis, while at the same time show strongly improved clinical response to anti-pyrimidine drugs. This result was further verified in another clinical dataset from the CTR-DB database, which contains sensitivity (response or no-response) to fluorouracil based chemotherapy^22^. We still observe significant enrichment of response patients in DeepMeta predicted high dependent group compared with low dependent group (**Figure S14C**).

### Metabolic vulnerabilities of cancers with undruggable driver genes

We considered three currently difficult-to-drug cancer driver genes, including *MYC*, *TP53* and *CTNNB1*. For each driver gene, we first screened metabolic enzymes whose predicted dependencies were correlated with the mutational status of the driver gene at the pan-cancer level (logistic regression Odds Ratio > 1 and *P*-value < 0.05, **Figure S15A**). Subsequently, in each cancer type, for each screened metabolic enzyme, we evaluated the dependency difference of the metabolic enzyme gene in samples with functional mutations in the driver gene compared to samples without the driver gene mutations, using a one-tailed Fisher’s test. The results were summarized in **Figure 4A and Figure S15B**. Several of these DeepMeta predicted metabolic vulnerabilities for cancer driver genes has been recapitulated in published literatures, such as CAD, TYMS for *MYC* overexpression, GPX4 for *TP53* inactivation^3, 6, 23, 24^ (**Figure S15A**). We have compiled a list summarizing the metabolic genes upon which tumor cells harboring functional mutations in driver genes such as *MYC*, *TP53*, and *CTNNB1* are dependent (**Figure S15A and B**).

**Figure 4:**
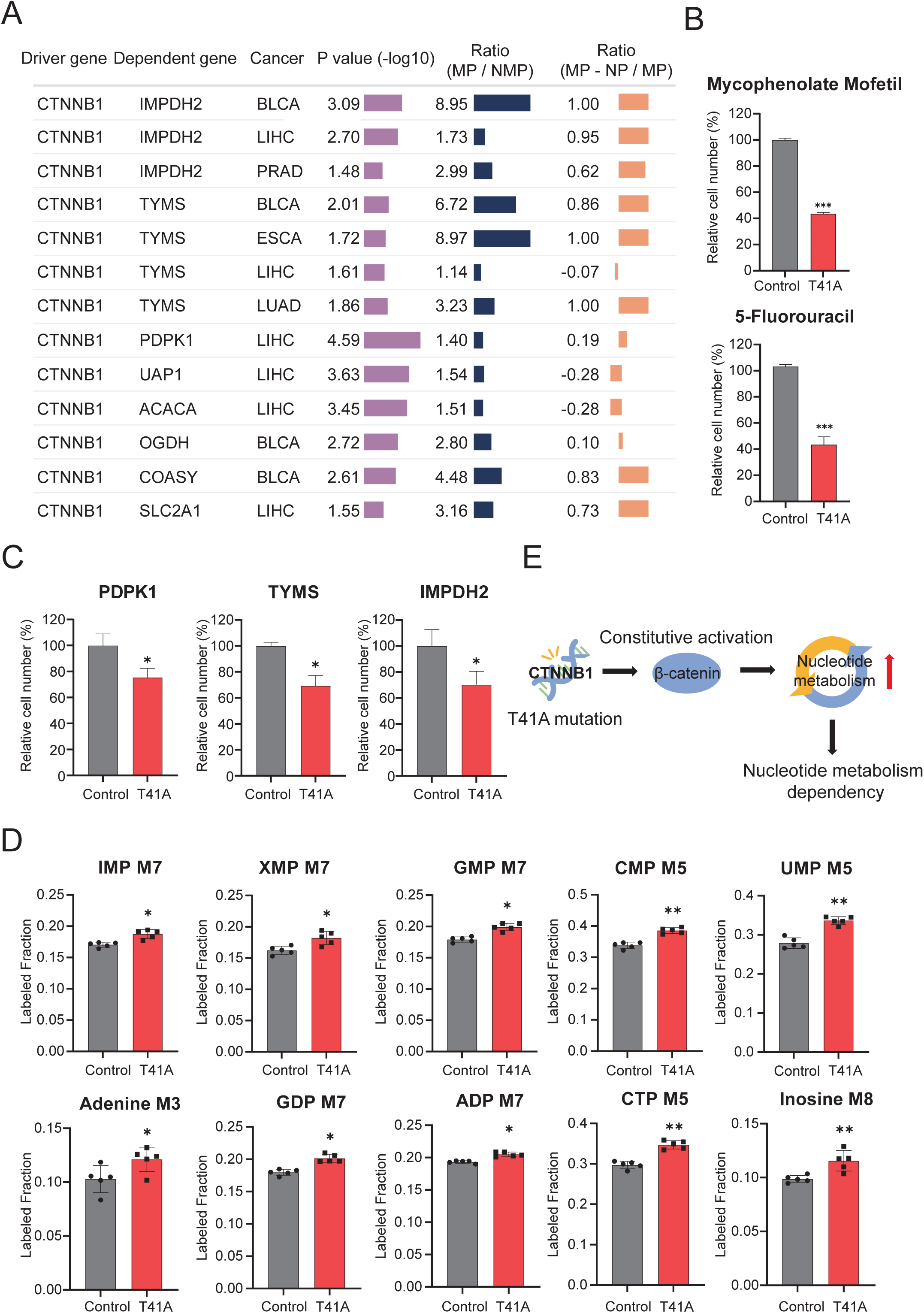
Experimentally validated metabolic targets for “undruggable” *CTNNB1* activating mutation harboring cells. **A,** Metabolic target screen results of *CTNNB1*. *P* values are calculated by one tail Fisher test. MP: Proportion of predicted positive cases in cancer samples with the indicated driver mutation, NMP: Proportion of predicted positive cases in cancer samples without the indicated driver mutation, NP: Proportion of predicted positive cases in corresponding normal tissues. **B,** Growth suppression of WI38 cells with *CTNNB1*-T41A mutation following treatment with mycophenolate mofetil (IMPDH2 inhibitor, 10uM) and 5-fluorouracil (TYMS inhibitor, 20uM). **C**, shRNA-mediated knockdown of *PDPK1, TYMS*, and *IMPDH2* leads to growth inhibition in *CTNNB1*-T41A mutant WI38 cell lines. **D**, Differential abundance of 13C-glucose-derived nucleotide metabolic intermediates between control and *CTNNB1*-T41A mutant WI38 cells. **E**, Schematic representation of mechanisms by which *CTNNB1* activating mutation harboring cells are more dependent on nucleotide metabolism and can be selectively inhibited by targeting nucleotide metabolism pathway. IMP, inosine monophosphate; XMP, xanthosine monophosphate; GMP, guanosine-5-monophosphate; CMP, cytidine 5-monophosphate; UMP, uridine 5-monophosphate; GDP, guanosine-5-diphosphate; ADP, adenosine-5-diphosphate; CTP, cytidine triphosphate; M+n, native metabolite mass (M) plus number of isotopically labeled carbons (n). Control: WI38 cells infected with a lentivirus carrying the vehicle control; T41A: WI38 cells stably expressing the CTNNB1-T41A mutation.

### Experimental validations of DeepMeta predicted metabolic dependencies

Building upon the predicted metabolic dependency genes delineated by DeepMeta, *CTNNB1* activating mutations leads to increased dependencies on nucleotide metabolism pathway (**Figure 4A**). We established stable cell lines harboring the *CTNNB1*-T41A mutation in WI38 and 293T cell lines. The drugs mycophenolate mofetil and 5-fluorouracil, known to target *IMPDH2* and *TYMS* respectively, were selected for further study. Significant selectivity in growth inhibition was exhibited by both mycophenolate mofetil and 5-fluorouracil in the *CTNNB1*-T41A mutant WI38 and 293T cell lines (**Figure 4B, Figure S16A**). In contrast, drugs targeting other molecules did not exhibit specific growth suppressive effects on *CTNNB1*-T41A mutant cell lines (**Figure S16C-D**). Subsequently, shRNA-mediated knockdown of *PDPK1*, *TYMS*, and *IMPDH2* also revealed a selective inhibitory effect on the growth of the *CTNNB1*-T41A mutant WI38 and 293T cell lines (**Figure 4C and Figure S16B**). It is well-documented that *IMPDH2* and *TYMS* are involved in the nucleotide metabolism pathways (**Figure S16E**). *AXIN1* is a crucial negative regulator in the Wnt/β-Catenin pathway and its knockout results in the sustained accumulation of β-Catenin. Our findings revealed that both 5-fluorouracil and mycophenolate mofetil exhibited selective inhibition of tumor cells with *AXIN1* knockout (**Figure S17A-B**). Subsequently, we utilized shRNA to specifically downregulate *IMPDH2*, *PDPK1*, and *TYMS* in control and *AXIN1* knockout MHCC97-H cells. The results, depicted in **Figure S17 C-E**, reveal that the shRNA-mediated downregulation of these genes significantly attenuates the proliferation of *AXIN1*-null cells. To probe the metabolic pathway alterations in nucleotide metabolism, we conducted non-targeted metabolomics studies using 13C-glucose isotope tracing to investigate the metabolic alterations in WI38 cells harboring the *CTNNB1*-T41A mutation. The results indicated an increased flux through nucleotide metabolic pathways in the *CTNNB1*-T41A WI38 cells (**Figure 4D**), suggesting a heightened dependency on nucleotide metabolism pathways, which could account for the selective growth inhibition observed when targeting these pathways (**Figure 4E**). Further investigations into other metabolic dependency targets predicted by our model were also conducted. We engineered A549 cell lines overexpressing gene *MYC* and harboring gene *TP53* mutation. SiRNA knockdown experiments confirmed the dependency of the *MYC* overexpressed, *TP53*-R273H mutant cell lines on the predicted metabolic targets (**Figure S18**). In summary, through *in vitro* experiments, we have validated the reliability of our model’s predictions and have identified novel metabolic vulnerability targets which could potentially be exploited for therapeutic interventions in cancer with “undruggable” driving alterations. These targets include, but are not limited to, key enzymes within the nucleotide synthesis pathways, which our data suggest may serve as pivotal checkpoints in the metabolic networks of cancer cells harboring specific driver mutations.

## Discussion

Many cancer are driven by genetic alterations that are not druggable on their own, for example *CTNNB1, MYC, TP53* etc. Metabolic enzymes or pathways are druggable targets for many kinds of diseases, including cancer. Cancer cells show significant heterogeneity in driving genetic alterations, and identify the metabolic vulnerability of each specific cancer cell is essential for applying metabolic targeting approaches in specifically killing cancer cells but not impairing normal tissue cells. Here we present a first systematic exploration of the precise metabolic dependency of cancer.

Metabolic vulnerabilities are long-sought targets for cancer treatment. Antimetabolites are the most widely used group of anticancer drugs. The first anti-cancer metabolism targeting drug aminopterin was developed in 1947^25^. As a folate analogue, Aminopterin inhibits one-carbon transfer reactions required for de novo nucleotide synthesis. Successful antimetabolites include purine analogues 6-mercaptopurine (6-MP) and 6-thioguanine (6-TG) and pyrimidine analogue 5-fluorouracil (5-FU), which inhibits purine and pyrimidine synthesis^26^. Based on this study, nucleotide metabolism shows pan-cancer dependency, and this is consistent with the wide range application of anti-nucleotide metabolites in cancer treatment. DeepMeta can be applied to identify the cancer patients who are particularly suitable for these antimetabolites treatment.

In summary, DeepMeta is unique in its ability to predict the metabolic dependency for individual cancer samples, could directly promote personalized precision medicine for cancer. DeepMeta incorporates two modules, one is sample status information represented by gene expression and the other is the metabolic enzyme network information, and the ablation analysis suggests the importance of both parts for the model’s performance. Thus, when modeling biological problems, incorporating prior knowledge into the model or data helps the model learn. In terms of metabolic dependency prediction, the metabolic network composed of metabolic genes is an important prior information. The performance of DeepMeta has been extensively validated in independent datasets. In TCGA samples, DeepMeta successfully identify the patients showing robust clinical response to anti-pyrimidine drugs. DeepMeta predicts the metabolic targets for cancers driven by “undruggable” genetic alterations, and these predictions can be experimentally validated. Further validation of in vivo model is needed in the future to better demonstrate the effectiveness of these predicted metabolic targets. The clinical application of DeepMeta will enable precision stratification and clinical management of cancer patients.

### Limitations of the study

DepMeta model was trained on cell line data and then applied on TCGA sample tissues. The expression of cultured clonal tumor cells could be different from the bulk tumor tissue of TCGA samples, and currently we still do not have tools to accurately extract tumor cell transcriptome from the bulk tumor tissue transcriptome. Although the analysis of taking cell type in tissue as a covariate (**Figure S13**) suggested that consider the transcriptome of bulk tumor tissue as a whole is practical, future advances in computational and experimental methods will allow for the more precise resolution of cancer metabolic dependencies.

## Resource availability

### Lead contact

Requests for further information and resources should be directed to and will be fulfilled by the lead contact, XueSong-Liu.

### Materials availability

This study did not generate new unique reagents. Data and code availability

All cell line CRISPR-Cas9 screen data and omics data used for analysis are publicly available from DepMap Data Portal at https://depmap.org/portal/. TCGA mRNA expression and mutation data are publicly available from Xena at https://xenabrowser.net, copy number data are available from GDC PanCanAtlas publication at https://gdc.cancer.gov/about-data/publications/<u>. All</u> <u>processed data, including DeepMeta prediction for cell line and TCGA are</u> <u>available in</u> https://github.com/XSLiuLab/DeepMeta.

The DeepMeta model, codes required to reproduce the analysis outlined in this manuscript are available in https://github.com/XSLiuLab/DeepMeta, and analysis report are available online in https://xsliulab.github.io/DeepMeta/.

## Acknowledgments

We thank ShanghaiTech University High Performance Computing Public Service Platform for computing services. We thank multi-omics facility, molecular and cell biology core facility of ShanghaiTech University for technical help. We thank Raymond Shuter for editing the text. This work is supported by Shanghai Science and Technology Commission (24J22800700), National Natural Science Foundation of China (82373149), cross disciplinary Research Fund of Shanghai Ninth People’s Hospital, Shanghai JiaoTong University School of Medicine (JYJC202227), open project fund of the National Health Commission’s key laboratory of individualized diagnosis and treatment of nasopharyngeal cancer (2023NPCCK02), startup funding from ShanghaiTech University. The graphical abstract was created by BioRender.

## Author contributions

TW developed the DeepMeta algorithm, performed the computational analysis and drafted the manuscript. XZ, YZ performed all the validation experiments. DQ, KD, DX, WW, XX, XL participated in critical project discussion and sharing of essential research materials. XSL conceptualized the idea, designed, supervised the study and wrote the manuscript.

## Supplemental information

Document S1. Figures S1-S18 and Tables S1 and S2

Table S3. Excel file containing additional data too large to fit in a PDF, Correspondence between cell line ID and GSM file.

## STAR Methods

### Experimental model and study participant details

Cell lines have been authenticated using DNA profiling using different and highly polymorphic short tandem repeat (STR) loci and were maintained in appropriate culture medium as suppliers suggested. A549 cells were incubated in RPMI-1640 medium (C11875500CP, Gibco) enriched with 10% fetal bovine serum (FBS, 10091148, Gibco) and penicillin/streptomycin (15140163, Gibco). AXIN1 knockout and sgRNA control MHCC97-H cell lines are a generous gift from Liye Zhang lab from Shanghaitech university. WI38 cells, MHCC97-H cells and HEK-293T cells were cultured in DMEM (C11995500CP, Gibco) supplemented with 10% FBS and penicillin/streptomycin. All cells were maintained at 37°C with 5% CO2.

## Method details

### Training and validation datasets

The training dataset used in this study was CRISPR-Cas9 screen of 17453 genes in 1078 cell lines from Depmap project^12^ (23Q2 version, https://depmap.org/portal/). We used the post-Chronos gene dependency probability score to set up a classification task for our model^27^. The dependency probabilities value is between 0 and 1. Cell line-gene pair with dependency score >0.5 were defined as the positive class and the cell line-gene pair with dependency score <=0.5 were defined as the negative class. We choose cancer cell lines with matched cancer type or tissue-specific Genome-Scale Metabolic Models (GSMs) in Metabolic Atlas database^28^ (**Table S3**). Genes existing in corresponding enzyme network and positive in at least 3 cell lines were chosen All together, 343,029 (760 cell lines and 1063 genes) labeled samples were available for training the model. We included other four independent validation datasets for model performance verification: 1) Another CRISPR screen data conducted by the Sanger institute using a different CRISPR library^14^; 2) RNAi screen data, which apply DEMETER2 to the combination of three large-scale RNAi screening datasets^15^: the Broad institute project Achilles, Novartis project DRIVE^16^, and the Marcotte et al. breast cell line dataset^17^ ; 3) The Childhood Cancer Model Atlas (CCMA) dataset which performs CRISPR-Cas9 screen in 110 paediatric solid tumour cell lines^18^. This dataset uses Z score to quantify gene dependency. Consistent with the original paper, we classified cell-gene pair with Z score less than -1.5 as positive class and others as negative class. 4) Pemovska et al drug screen dataset^19^, which performs metabolic drug library (contains 243 compounds) screen in 15 cancer cell lines and reports AUC scores. TCGA drug treatment data was curated by Ding. et.al^29^ and was downloaded from http://lifeome.net/supp/drug_response/. 5) PRISM repurposing dataset, which contains the results of pooled-cell line chemical-perturbation viability screens for 4,518 compounds screened against 578 cell lines and the drug effect score is represented by log fold change (cell viability between treatment and control).

### Omics dataset

The gene expression data of 19193 genes in 1408 cell line was download from Depmap portal (23Q2 version). The gene expression data of normal samples was download from GTEx^30^ portal (https://www.gtexportal.org/) and we used median gene-level data for each tissue type. The gene expression level was represented as TPM (transcripts per million). Overall, we chose 7993 genes with standard deviation (SD) > 1 in tumor samples or normal tissues. We defined the cell gene differential expression vector as:

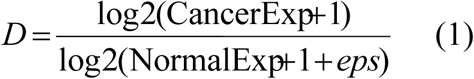

Where eps is used to avoid the situation where the denominator is 0, resulting in infinity, the eps value we used here is 0.01.

For TCGA samples, gene expression in TPM unit and somatic mutation data generated by the multi-center mutation calling in multiple cancers (MC3) project^31^ were download from Xena^32^ (https://xenabrowser.net). GISTIC2.0^33^ gene-level focal copy number score file was downloaded from GDC PanCanAtlas publication (https://gdc.cancer.gov/about-data/publications/PanCan-MYC-2018).

### Metabolic network dataset

Cancer-specific or tissue-specific genome-scale metabolic models (GSMs) were download from metabolic atlas repository (http://www.metabolicatlas.org/) in the compressed systems biology markup language (SBML) format. The cancer-specific metabolic network was constructed by INIT algorithm, as described in Rasmus Agren et.al^34^. For a cell line, if there is a cancer-specific GSM corresponding to its cancer type, then we use that GSM, otherwise we use a GSM from the same tissue origin as the cell line. Then we applied the R package Met2Graph^35^ to extract enzyme network from GSMs. In this network, enzymes are the nodes connected by edges represented by metabolites. Two enzymes are connected if they catalyze two reactions which produce or consume a specific metabolite.

### Pathway dataset

To obtain the features of genes, we collected the chemical and genetic perturbations (CGP) gene sets from MSigDB database^11^. After filter gene sets without any gene existed in GSMs, we got 3247 gene sets. For each gene to be predicted, its feature is a 3247-dimensional 0/1 vector defined by:

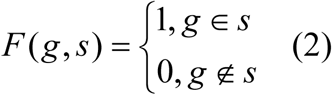

Where g and s donate specific gene and CPG gene set respectively.

For metabolic gene annotation, we filtered metabolism pathway class from KEGG^36^ (https://www.genome.jp/kegg/) database and resulted in 87 metabolism pathway containing 1788 genes.

### Deep learning model design

We present the DeepMeta framework to predict the metabolic gene dependence based on the metabolic pathway information characterized by enzyme networks and sample status information defined by gene expression. Thus, DeepMeta has two inputs, the sample-specific enzyme network and the gene expression information. The sample-specific enzyme network (SEN) is constructed from cancer-specific or tissue-specific GSMs, and in this network, enzymes are the nodes connected by edges represented by metabolites. SEN are constructed by removing gene nodes in the cancer-specific enzyme network that are not expressed in the sample (defined as expression level below 1 TPM) and corresponding edges. The features of gene nodes in the enzyme network are composed of 3247-dimensional binary (0/1) vector, indicating whether the gene exist or not in the collected 3247 CPG gene sets. Thus different gene nodes would have different features. The gene expression information was represented by expression of genes in cancer relative to normal tissue, which reflects dysregulated cancer cell state. We choose 7993 genes with SD of expression > 1 TPM in tumor cell or normal tissues. DeepMeta has two functional modules to obtain metabolic enzyme and sample state representations, and get the final enzyme dependency prediction by combining these two representations (**Figure 1A**). DeepMeta uses graph attention network (GAT) to get gene embedding from SEN *S* . GAT allows gene nodes to aggregate information from their neighborhoods, with self-attention mechanisms, thus learn the relative importance of each neighbor during the aggregation process. For a given node *i*, the embedding of all its neighbor nodes *j* in layer *l*, 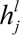 are used to compute embedding of enzyme gene node *i* on the next layer 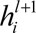 gene feature *F*):, using the following equation (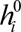 is the original

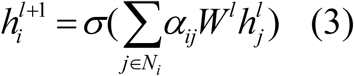

Where *W* is the matrix of learnable weights of layer *l*, *N _i_* is the set of *i ’ s* neighbors and includes *i* itself, *σ* is the non-linear activation function and we used RELU here. *α_ij_* is the learnable attention weight between node *i* and *j*, it can be obtained from normalizing attention coefficient *e_ij_* by softmax function where attention coefficient was calculated by attention mechanism *a* :

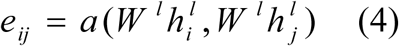

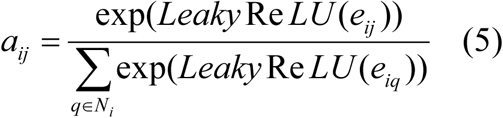

We applied multi-head attention mechanisms to benefit robust node representations. We concatenated the *K* independent attention heads to get the output gene embedding of this layer:

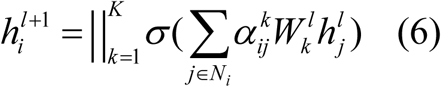

Where || is concatenation operation, 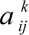 is 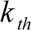 th normalized attention weight.

Thus, for one specific gene node *i*, we can get its final gene embedding *G_i_* after several GAT layers of message passing transformation from its original feature vector *F* . We utilized a three-layer GAT architecture here because deeper graph convolutional networks are prone to over-smoothing^37^.

For one specific cell line or sample *^m^*, the sample differential expression vector *^D^* was transformed to sample embedding *C* by multi-layer perceptron (dense layers):

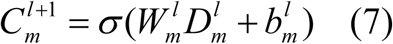

Where *^Wl^* is learnable weights of dense layer *l*, *^bl^* is learnable bias of dense layer *l* .

For metabolic dependency prediction, we concatenated the embedding of sample-gene pairs and transformed them by external dense layers into one output probability nodes to predict the dependency, i.e.

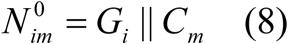

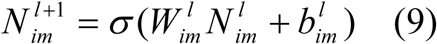

Where *N* is the representation vector of the intermediate layer and contains one value in the final output layer 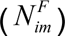. After Sigmoid function transformation, we can get value between 0 and 1, representing dependency probability:

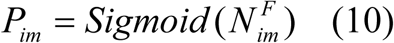

The object of DeepMeta is to optimize the binary cross-entropy (BCE) loss:

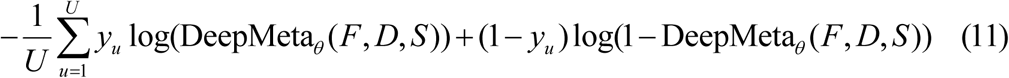

Where *U* is number of sample-gene pairs in one mini-batch, *θ* is parameters of the model DeepMeta, *^F^*, *^D^* and *S* are enzyme gene features, sample differential expression vector and enzyme network respectively.

### Model training and evaluation

The entire model was trained using the data from CRISPR-Cas9 knockout screens of 1063 enzyme gene in 760 cell lines (in total, 343,029 samples). We used random sampling 90% (307,166 samples) to train the model, 10% (35,863 samples) to test model performance, 30% of training data to monitor training process and tune hyper-parameters. The number of layers of sample encoding module and the final fully connected layers are both three. After grid-searching hyper-parameter tuning, we set following hyper-parameters for DeepMeta: 512 for hidden layer dimensions of GAT, 3 for number of heads of GAT, 4000, 1500, 512 for hidden layer dimensions of sample state encoder, 1024, 512 for hidden layer dimension of final fully connected layer module, learning rate 0.001, and training 15 epochs. We used common classification model performance evaluation metrics, including the area under the receiver operating characteristic curve (AUROC), the area under the precision-recall curve (AUPRC), and the F1 value. The ratio of negative to positive samples in our dataset is 3:1, thus we also used Kappa value, which is suitable for imbalanced data.

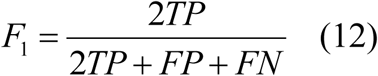

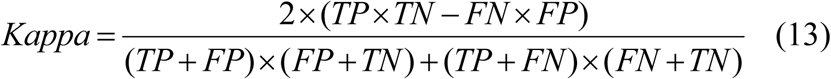

Where TP is for true positive, FP is for false positive, FN is for false negative, and TN is for true negative.

We also evaluated the prediction performance for individual genes by AUROC (termed per-ROC). We focused on 314 diverse genes whose proportion of cell lines with positive labels ranged from 10% to 90%.

### Model comparison

We performed ten-fold cross-validation analyses to compare model performance. Cell lines were randomly partitioned into ten equal-size folds. For one training process, a single fold was retained as the validation data for testing the model, and the remaining nine folds were used as training data. The cross-validation process was then repeated ten times, with each of the ten folds used exactly once as the validation data. We calculated and compared AUROC, AUPRC, F1, Kappa and per-ROC score for these validation data across different models.

In ablation experiments, for metabolic gene features, which are represented by a binary vector with relatively few 1s, so we changed all 1s into 0s, then randomly sampled the same number of 0s and converted them into 1s, forming a random gene feature vector for model training. Regarding sample expression, the gene expression of sample was shuffled randomly to create a randomized sample expression matrix for model training. In addition, we compared DeepMeta with the model that do not utilize enzyme network as input to demonstrate the advantages of GAT in extracting information. In this scenario, we combined gene features of the original metabolic network (3247-dimensional) and sample expression profile features (7993-dimensional) to form the new gene feature (a total of 11240 dimensions), and then used a simple four-layer MLP to process this feature to predict the metabolic vulnerability of the gene. We also performed label scrambling analysis to show that the model performance we obtained is not due to random chance. We randomly permuted the sample labels to generate mismatched input and output data, and used these data for model training and performance validation.

The performance of DeepMeta was compared to several machine learning models. We used three classic machine learning algorithms, including logistic regression, random forest, and support vector machines (SVM). The input data of these three models are consistent with DeepMeta (gene node features and gene expression features). Since these machine learning models cannot be extended to high-dimensional input data, we first performed dimensionality reduction using PCA on the input data, and retained the top 100 PC dimensions for gene features and cell expression respectively (a total of 200 dimensions). The reduced features are fed into the machine learning model for training and performance validation. All machine learning models were implemented using corresponding functions in scikit-learn^38^ python library. Logistic regression model was implemented by “SGDClassifier” function with loss of log-loss. Random forest was implemented by “RandomForestClassifier” function. The SVM contains linear and RBF (radial basis function) kernel, being implemented by “SGDClassifier” function with loss parameter of hinge and Nystroem method for kernel approximation, respectively. SVM with nonlinear radial basis function kernels did not converge and was not shown in results.

DeepDep is transfer learning based model for predicting gene dependency^21^, we used the independent CCMA dataset^18^ and drug screening dataset^19^ to compare the performance of the DeepDep with our model DeepMeta. We first used the R package “Prep4DeepDEP” developed by the author to process the input data, using only gene expression as input, and the predicted genes list were consistent with those predicted by our model. Then applied the Python script (PredictNewSamples.py) provided by the author to make predictions (the DeepDep model file used was model_exp_paper.h5). The value DeepDep predicted is not probability value, but gene effect scores. Therefore, according to the instructions on the DepMap official website, samples with predicted values less than -0.5 are labeled as positive examples, and others as negative examples and then compare differences of CCMA Z score between the two categories using Wilcoxon test.

### Model interpretation

We interpreted the prediction results of the model from the node neighbor perspectives. We used the attention weights of graph attention model to get the importance of a gene’s neighbor genes for its prediction (**Figure S6A**). We extract neighbor genes with top 3 attention weight for gene A in each true positive and true negative cell lines (attention weight is the average of 3 attention heads). We define the neighbor importance score of gene B to gene A as the frequency of gene B in extracted neighbor genes of gene A divided by the number of cell lines where gene A is exist. The visualization was carried out using the R package “igraph”^3939^, wherein the weights of the edges correspond to the neighbor importance scores.

### TCGA analysis

We applied DeepMeta to the TCGA dataset, consisting of 25 cancer types and 8937 samples. The construction of sample differential expression matrix and sample-specific metabolic enzyme network is the same as described above.

For each sample, in order to calculate pathway level dependency, we employed a one-tailed Fisher’s exact test to assess the enrichment of genes predicted to be positive within a specific metabolic pathway relative to all other pathways in that sample. The odds ratio (OR) and *P* value were calculated to quantify the strength of this enrichment. The OR is defined as the ratio of the odds of positive gene enrichment in the target pathway to the odds of positive gene enrichment in all other pathways. Mathematically, the OR is given by:

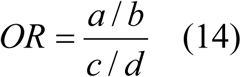

Where a is the number of genes predicted to be positive in the target pathway, b is the number of genes predicted to be negative in the target pathway, c is the number of genes predicted to be positive in all other pathways combined, d is the number of genes predicted to be negative in all other pathways combined.

To mitigate the impact of pathway gene set size on enrichment results, we also conducted a permutation test. For a given pathway in a particular sample, we randomly sampled genes equal to the number of genes in that pathway to form a “random gene set”. We calculated the proportion of genes predicted as positive in this random gene set, repeated this process 1000 times to obtain 1000 proportion values. Then, we compared this empirical distribution of 1000 values with the actual proportion of positive genes in the pathway to obtain a *P*-value. Thus, for each sample, we can get such a permutation test *P* value:

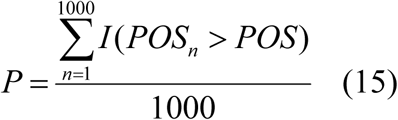

Where *POS*_n_ is the proportion of genes predicted as positive in random gene set of one permutation, *POS* is the proportion of genes predicted as positive in the actual pathway gene set. *I* is an indicator function.

Kaplan–Meier survival analysis was performed using the R package “survival” with log-rank test, and Cox proportional hazard analysis was performed using the R package “ezcox”. In all survival analyses, we filtered for cancer types with a sample size greater than 10. All cancer types were mixed together (we also show the results by cancer type) in K-M survival analysis, while in Cox analysis we considered cancer type as a control variable to account for the effect of cancer types.

In cell cycle signature analysis, we considered 446 cell cycle related genes collected by Lundberg et.al^40^. Gene set variation analysis^41^ (GSVA) was applied in TCGA expression data (input as log2TPM) to calculate enrichment score of the cell cycle gene set. This procedure was implemented by R package “GSVA”.

### Metabolic target prediction for undruggable cancer driver genes

We applied DeepMeta in TCGA dataset to predict metabolic targets for some important undruggable cancer genes, including *MYC* (*MYC*, *MYCL1*, *MYCN*), *TP53* and *CTNNB1*. For the *MYC* gene, as indicated in Schaub et al^42^, a corrected value > 0 of focal copy number variation was considered as amplification. For *TP53* and *CTNNB1*, we annotated gene mutations using variant information (loss of function for *TP53* or gain of function for *CTNNB1*) existed in The Clinical Knowledgebase^43^ (CKB) database. For a specific gene, the presence of aforementioned amplification or functional mutations in a sample designated the sample as “mut-sample”, while samples lacking any mutations in the gene were labeled as “non-mut samples”. To screen potential metabolic targets of these cancer genes, we utilized logistic regression to model the predicted dependency label of metabolic genes and the mutational status of the cancer gene, including cancer type as covariates to account for the effect of cancer types, and reporting corresponding odds ratio (OR) and *P* values of mutation status variable:

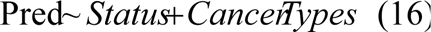

For a specific metabolic gene, the OR value of the mutation status variable is the ratio of the proportion of sample predicted to be dependent on this metabolic gene in the “mut-sample” and proportion of sample predicted to be dependent on this metabolic gene in the “non-mut sample”.

After screening for significant metabolic genes in pan-cancer level (logistic regression *P* < 0.05 and OR > 1), we applied one tail fisher test to analyzed these genes per cancer type. In more details, within each cancer type, we applied one-tailed Fisher’s exact test to check whether samples predicted to be dependent on specific metabolic gene are enriched in mutated samples, compared to non-mutated samples and report corresponding *P* values.

### Generation of mutant expressing cell lines

pCDH-CMV-MCS-EF1-Puro plasmid was digested with Xho I (1635, TaKaRa) and BamH I (1605, TaKaRa) restriction endonuclease followed by purification from an agarose gel using HiPure gel Pure DNA extraction kit (D2110-02, Magen). The mutant genes, *CTNNB1* T41A, MYC, TP53-R273C, were then ligated into the linearized plasmid vector using T4 DNA ligase (M0202S, NEB) to construct pCDH-mutant genes recombinant plasmids. On day 1, plate HEK-293T cells in a 6 cm culture dish with 5 mL of DMEM supplemented with 10% FBS, without antibiotics, and incubated at 37°C with 5% CO2. On the following day, prepared the transfection complex using OPTI-MEM reduced serum medium which contained 1 μg of the pCDH-mutant genes recombinant plasmids, 750 ng of the psPAX2 packaging plasmid, 250 ng of the pMD2.G envelope plasmid. According to the manufacturer’s instructions, using OPTI-MEM reduced serum medium dilute transfection reagent EZ Trans (LIFE iLAB BIO, Shanghai, China). The two components were mixed by gentle pipetting for 3-5 times and incubated at room temperature for 20 minutes. Slowly add the mixture along the inner wall of the cell culture dish into the culture medium. After 8 hours post-transfection, 5 mL of fresh antibiotic-containing DMEM was added to the cells. Following a 48-hour incubation period post-transfection, the medium was harvested and passed through a 0.45 μm filter to remove cellular debris. The virus-containing medium could be temporarily stored at 4°C. Target cells, WI38, HEK-293T cells and A549 cells, were seeded at an appropriate density to reach 20% confluence on the day of transduction. The cells were incubated with the lentivirus-containing medium for a duration of 48 hours. Subsequently, the medium was discarded, and cells were replenished with fresh complete culture medium. After an overnight incubation, a selective medium containing puromycin (ST551-10mg, beyotime) at a final concentration of 2 µg/mL was applied to the cultures. The cells were subjected to puromycin selection for a defined period to enrich for a stably transduced population expressing the gene of interest.

### Generation of shRNA knockdown cell lines

To commence the gene-silencing study, short hairpin RNA (shRNA) oligonucleotides targeting *IMPDH2* (target sequence: CCACAACTGTACACCTGAA) and *TYMS* (target sequence: CAATCCGCATCCAACTATT) were synthesized. These oligonucleotides were subsequently cloned into the pLKO.1-GFP plasmid using standard molecular biology techniques. The pLKO.1 plasmid was digested with AgeI (R3552S, NEB) and EcoRI (R3101V, NEB) restriction enzymes, followed by purification from an agarose gel using HiPure gel Pure DNA extraction kit (D2110-02, Magen). The annealed oligonucleotides were then ligated into the linearized plasmid vector using T4 DNA ligase (M0202S, NEB). Transformation of the ligation product was performed using competent DH5α E.coli cells, and clones were identified by DNA sequencing. For the generation of stable knockdown cell lines, lentiviral particles were produced in HEK-293T cells. On day 1, HEK-293T cells were plated in a 6 cm culture dish with 5 mL of DMEM supplemented with 10% FBS, without antibiotics, and incubated at 37°C with 5% CO2. On the following day, transfection was carried out in the late afternoon using an optimized polypropylene microfuge tube protocol. The transfection mixture contained 1 μg of the pLKO.1-shRNA plasmid, 750 ng of the psPAX2 packaging plasmid, 250 ng of the pMD2.G envelope plasmid, and OPTI-MEM reduced serum medium. The proprietary transfection reagent EZ Trans (LIFE iLAB BIO, Shanghai, China) was used according to the manufacturer’s instructions. After 7 hours post-transfection, 5 mL of fresh antibiotic-containing DMEM was added to the cells. Following a 48-hour incubation period post-transfection, the medium was harvested and passed through a 0.45 μm filter to remove cellular debris. The virus-containing medium could be temporarily stored at 4°C. For viral transduction, target cells were plated in 6 cm plate and incubated overnight at 37°C with 5% CO2, aiming for approximately 20% confluency. Lentiviral particles were then added to each plate. At 48 hours post-infection, fresh medium was replenished and cells were further cultured overnight. The successfully transduced cells expressing GFP were isolated utilizing flow cytometry-based cell sorting.

### siRNA knockdown of target genes

Target cells, A549 cells, with overexpression *MYC* gene and mutant genes *TP53* R273C were plated into 6-well plates at a density of 2×104 cells per well, with 2.5 mL of DMEM supplemented with 10% FBS, without antibiotics, and incubated at 37°C with 5% CO2. Target cells were transfected with 50 pmol of siRNA targeting the gene of interest (**Table S1**), or a non-targeting control siRNA (si-ctrl), utilizing the EZ-Trans transfection reagent according to the manufacturer’s protocol. Forty-eight hours post-transfection, the cells were harvested, and the silencing efficiency was quantitatively assessed by RT-qPCR to determine the expression levels of the target gene.

### Cell viability assay

In vitro inhibitor studies were conducted with compounds initially solubilized in DMSO to create a 10 mM stock solution, which was subsequently aliquoted and stored at -20°C. Detailed information regarding the compounds utilized is available in the Supplementary Material (**Table S2**). For proliferation assays, both control cells and cells engineered to express mutant genes or AXIN1 knockout cells were seeded into 96-well plates at a density of 4,000 cells per well. Triplicate wells were assigned for each condition within the control and experimental groups. Growth media containing various concentrations of the compounds were prepared, with the vehicle control comprising growth medium supplemented solely with DMSO. After a 72-hour incubation period, cell proliferation was assessed using the Cell Counting Kit-8 (CCK-8, C0005, TargetMol) according to the manufacturer’s instructions. All experimental conditions, including the control, were normalized to the cell proliferation observed in wells treated with growth medium supplemented only with DMSO.

For gene knockdown experiments, control cells and those expressing mutant genes were transfected with siRNAs targeting the predicted genes. Post transfection, cells were cultured for an additional 72 hours before evaluation of cell viability and proliferation using the CCK-8 assay, performed as specified by the provided protocol.

### Stable-isotope tracing experiments

In order to conduct stable isotope tracking experiments, the following protocols were strictly followed. 24 hours prior to isotope labeling, the culture medium was aspirated from the dishes and replaced with unlabeled fresh complete growth medium. Two hours before the incorporation of the stable isotope, the medium was once again replaced with fresh culture medium. Upon reaching the desired confluency of 80%, the culture medium was carefully aspirated, and cells were washed twice with 1X PBS to remove unlabeled medium. The culture medium was replaced as glucose-free DMEM (11966, Gibco) containing 4.5 mg/mL U-13C-glucose and 10% dialyzed fetal bovine serum (dFBS). Metabolite extraction solvent was prepared using a methanol/water mixture (4/1, v/v) and pre-chilled at -80°C for one hour prior to extraction to ensure it was ice-free. After 24 hours of labeling, the culture medium was completely removed and cells were washed twice with 2 ml 1× PBS. The culture dishes were placed on dry ice, and 1000 µl of cold metabolite extraction solvent was added per dish. The dishes were then stored at -80°C for at least 40 minutes. Following the extraction, the entire contents of the dish were scraped and transferred into 1.5 ml microcentrifuge tubes. The samples were vortexed for 1 minute at 4°C. Centrifugation was performed at 13,000 rpm for 10 minutes at 4°C to pellet insoluble material. The supernatant was carefully transferred to a new 1.5 ml microcentrifuge tube and dried using a vacuum concentrator at 4°C. The dried samples were stored at -80°C until further transport to the LC-MS for analysis.

### LC-MS analysis and data processing

LC–MS analysis was performed using a Waters ACQUITY UPLC I-Class system (Waters) coupled with a quadruple time-of-flight mass spectrometer (TripleTOF 6600, SCIEX). The mobile phases consisted of 25 mM ammonium acetate and 25 mM ammonium hydroxide in 100% water (mobile phase A) and 100% acetonitrile (mobile phase B). The instrumental parameters were optimized according to the methodologies previously reported^44^. Raw data acquired in .wiff format from the mass spectrometer were converted into .mzXML and .mgf files using the proteowizard tool MSConvert (version 3.0.23109). Then, according to the method mentioned above^44^, generate MS1 peak table and MS2 files, and submit them to the MetDNA^45^ (http://metdna.zhulab.cn/) for metabolite annotation. MetTracer (https://github.com/ZhuMetLab/MetTracer/) was employed for the detection of metabolite isotopologues within the nucleotide metabolism pathways.

### Quantification and statistical analysis

Statistical analysis of *in vitro* experiments was performed using GraphPad Software. All quantitative data are presented as the mean ± SD. Data in the bar graphs represent the fold change or the percentage relative to control with the SD of 3 independent experiments. Student’s t test was performed to compare 2 groups of independent samples. A *P* value of less than 0.05 was considered statistically significant (\**P* <0.05, ** *P* <0.01, *** *P* <0.001 in the Figure S). Other statistical details of each experiment can be found in the figure legend or the STAR Methods section.

## References

1. Hanahan D. Hallmarks of Cancer: New Dimensions. Cancer Discov 12, 31–46 (2022).

2. Duffy MJ, Crown J. Drugging “undruggable” genes for cancer treatment: Are we making progress? Int J Cancer 148, 8–17 (2021).

3. Liu T, et al. MYC predetermines the sensitivity of gastrointestinal cancer to antifolate drugs through regulating TYMS transcription. EBioMedicine 48, 289–300 (2019).

4. Kim J, et al. The hexosamine biosynthesis pathway is a targetable liability in KRAS/LKB1 mutant lung cancer. Nat Metab 2, 1401–1412 (2020).

5. Ying H, et al. Oncogenic Kras maintains pancreatic tumors through regulation of anabolic glucose metabolism. Cell 149, 656–670 (2012).

6. Tahaney WM, et al. Abstract GS1-09: Inhibition of GPX4 induces preferential death of p53-mutant triple-negative breast cancer cells. Cancer Research 82, GS1-09–GS01-09 (2022).

7. Li H, et al. CGMega: explainable graph neural network framework with attention mechanisms for cancer gene module dissection. Nat Commun 15, 5997 (2024).

8. Shu H, et al. Modeling gene regulatory networks using neural network architectures. Nat Comput Sci 1, 491–501 (2021).

9. Pacini C, et al. Integrated cross-study datasets of genetic dependencies in cancer. Nat Commun 12, 1661 (2021).

10. Veličković P, Cucurull G, Casanova A, Romero A, Lio P, Bengio Y. Graph attention networks. arXiv preprint arXiv:171010903, (2017).

11. Liberzon A, Subramanian A, Pinchback R, Thorvaldsdóttir H, Tamayo P, Mesirov JP. Molecular signatures database (MSigDB) 3.0. Bioinformatics 27, 1739–1740 (2011).

12. Tsherniak A, et al. Defining a Cancer Dependency Map. Cell 170, 564–576 e516 (2017).

13. Jia W, Sun M, Lian J, Hou S. Feature dimensionality reduction: a review. Complex & Intelligent Systems 8, 2663–2693 (2022).

14. Behan FM, et al. Prioritization of cancer therapeutic targets using CRISPR-Cas9 screens. Nature 568, 511–516 (2019).

15. McFarland JM, et al. Improved estimation of cancer dependencies from large-scale RNAi screens using model-based normalization and data integration. Nat Commun 9, 4610 (2018).

16. McDonald ER, 3rd, et al. Project DRIVE: A Compendium of Cancer Dependencies and Synthetic Lethal Relationships Uncovered by Large-Scale, Deep RNAi Screening. Cell 170, 577–592.e510 (2017).

17. Marcotte R, et al. Functional Genomic Landscape of Human Breast Cancer Drivers, Vulnerabilities, and Resistance. Cell 164, 293–309 (2016).

18. Sun CX, et al. Generation and multi-dimensional profiling of a childhood cancer cell line atlas defines new therapeutic opportunities. Cancer Cell 41, 660–677 e667 (2023).

19. Pemovska T, et al. Metabolic drug survey highlights cancer cell dependencies and vulnerabilities. Nat Commun 12, 7190 (2021).

20. Soula M, et al. Metabolic determinants of cancer cell sensitivity to canonical ferroptosis inducers. Nat Chem Biol 16, 1351–1360 (2020).

21. Chiu YC, et al. Predicting and characterizing a cancer dependency map of tumors with deep learning. Sci Adv 7, (2021).

22. Liu Z, et al. CTR-DB, an omnibus for patient-derived gene expression signatures correlated with cancer drug response. Nucleic Acids Research 50, D1184–D1199 (2022).

23. Miltenberger RJ, Sukow KA, Farnham PJ. An E-box-mediated increase in cad transcription at the G1/S-phase boundary is suppressed by inhibitory c-Myc mutants. Mol Cell Biol 15, 2527–2535 (1995).

24. Chong YC, et al. Targeted Inhibition of Purine Metabolism Is Effective in Suppressing Hepatocellular Carcinoma Progression. Hepatol Commun 4, 1362–1381 (2020).

25. Farber S, Diamond LK. Temporary remissions in acute leukemia in children produced by folic acid antagonist, 4-aminopteroyl-glutamic acid. N Engl J Med 238, 787–793 (1948).

26. Dey P, Kimmelman AC, DePinho RA. Metabolic Codependencies in the Tumor Microenvironment. Cancer Discov 11, 1067–1081 (2021).

27. Dempster JM, et al. Chronos: a cell population dynamics model of CRISPR experiments that improves inference of gene fitness effects. Genome Biol 22, 343 (2021).

28. Li F, Chen Y, Anton M, Nielsen J. GotEnzymes: an extensive database of enzyme parameter predictions. Nucleic Acids Res 51, D583–d586 (2023).

29. Ding Z, Zu S, Gu J. Evaluating the molecule-based prediction of clinical drug responses in cancer. Bioinformatics 32, 2891–2895 (2016).

30. The Genotype-Tissue Expression (GTEx) project. Nat Genet 45, 580–585 (2013).

31. Ellrott K, et al. Scalable Open Science Approach for Mutation Calling of Tumor Exomes Using Multiple Genomic Pipelines. Cell Syst 6, 271–281.e277 (2018).

32. Goldman MJ, et al. Visualizing and interpreting cancer genomics data via the Xena platform. Nat Biotechnol 38, 675–678 (2020).

33. Mermel CH, Schumacher SE, Hill B, Meyerson ML, Beroukhim R, Getz G. GISTIC2.0 facilitates sensitive and confident localization of the targets of focal somatic copy-number alteration in human cancers. Genome Biol 12, R41 (2011).

34. Agren R, Bordel S, Mardinoglu A, Pornputtapong N, Nookaew I, Nielsen J. Reconstruction of genome-scale active metabolic networks for 69 human cell types and 16 cancer types using INIT. PLoS Comput Biol 8, e1002518 (2012).

35. Granata I, Manipur I, Giordano M, Maddalena L, Guarracino MR. TumorMet: A repository of tumor metabolic networks derived from context-specific Genome-Scale Metabolic Models. Sci Data 9, 607 (2022).

36. Kanehisa M, Furumichi M, Tanabe M, Sato Y, Morishima K. KEGG: new perspectives on genomes, pathways, diseases and drugs. Nucleic Acids Res 45, D353–d361 (2017).

37. Li Q, Han Z, Wu X-M. Deeper insights into graph convolutional networks for semi-supervised learning. In: Proceedings of the AAAI conference on artificial intelligence) (2018).

38. Pedregosa F, et al. Scikit-learn: Machine learning in Python. the Journal of machine Learning research 12, 2825–2830 (2011).

39. Han W-S, Lee J, Pham M-D, Yu JX. iGraph: a framework for comparisons of disk-based graph indexing techniques. Proceedings of the VLDB Endowment 3, 449–459 (2010).

40. Lundberg A, Yi JJJ, Lindström LS, Tobin NP. Reclassifying tumour cell cycle activity in terms of its tissue of origin. NPJ Precis Oncol 6, 59 (2022).

41. Hänzelmann S, Castelo R, Guinney J. GSVA: gene set variation analysis for microarray and RNA-seq data. BMC Bioinformatics 14, 7 (2013).

42. Schaub FX, et al. Pan-cancer Alterations of the MYC Oncogene and Its Proximal Network across the Cancer Genome Atlas. Cell Syst 6, 282–300.e282 (2018).

43. Patterson SE, Statz CM, Yin T, Mockus SM. Utility of the JAX Clinical Knowledgebase in capture and assessment of complex genomic cancer data. NPJ Precis Oncol 3, 2 (2019).

44. Wang R, et al. Global stable-isotope tracing metabolomics reveals system-wide metabolic alternations in aging Drosophila. Nat Commun 13, 3518 (2022).

45. Zhou Z, Luo M, Zhang H, Yin Y, Cai Y, Zhu ZJ. Metabolite annotation from knowns to unknowns through knowledge-guided multi-layer metabolic networking. Nat Commun 13, 6656 (2022).

