## Supplementary material for "Precise metabolic dependencies of cancer through deep learning and validations": Figures S1-S18 and Tables S1 and S2

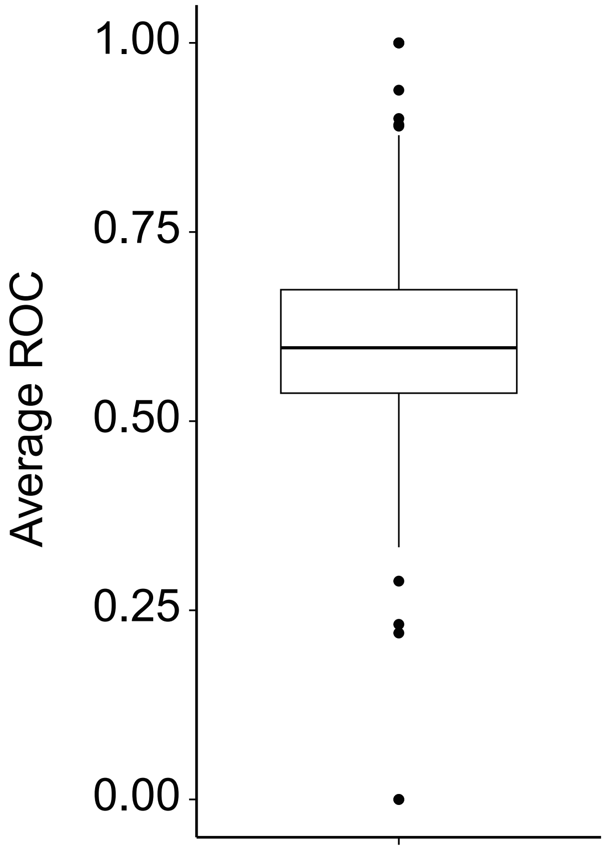


**Figure S1. The Distribution of ROC values for individual metabolic enzyme genes in ten-fold cross-validation.** We focused on 314 diverse genes whose proportion of cells with positive labels ranged from 10% to 90%. Values shown are average AUROC of ten test fold.

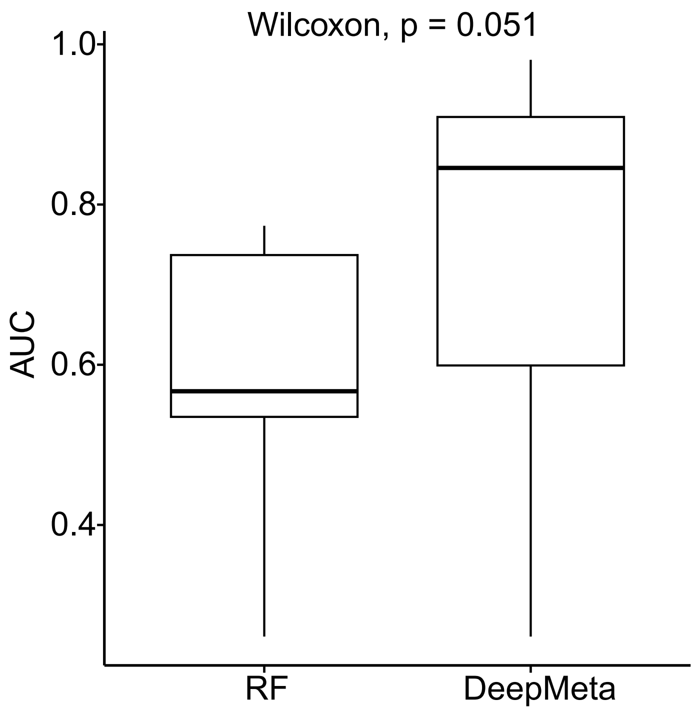


**Figure S2. Performance of random forest (RF) model on drug screening dataset and comparison with DeepMeta.** In each cell, the metabolic enzyme gene was assigned a rank value based on AUC score of drugs targeting this enzyme. We extracted the overlap genes between top 5 genes predicted by models (RF or DeepMeta) and the genes with top 3 AUC rank in each cell.Then the AUC scores of these overlap genes were compared between RF and DeepMeta.

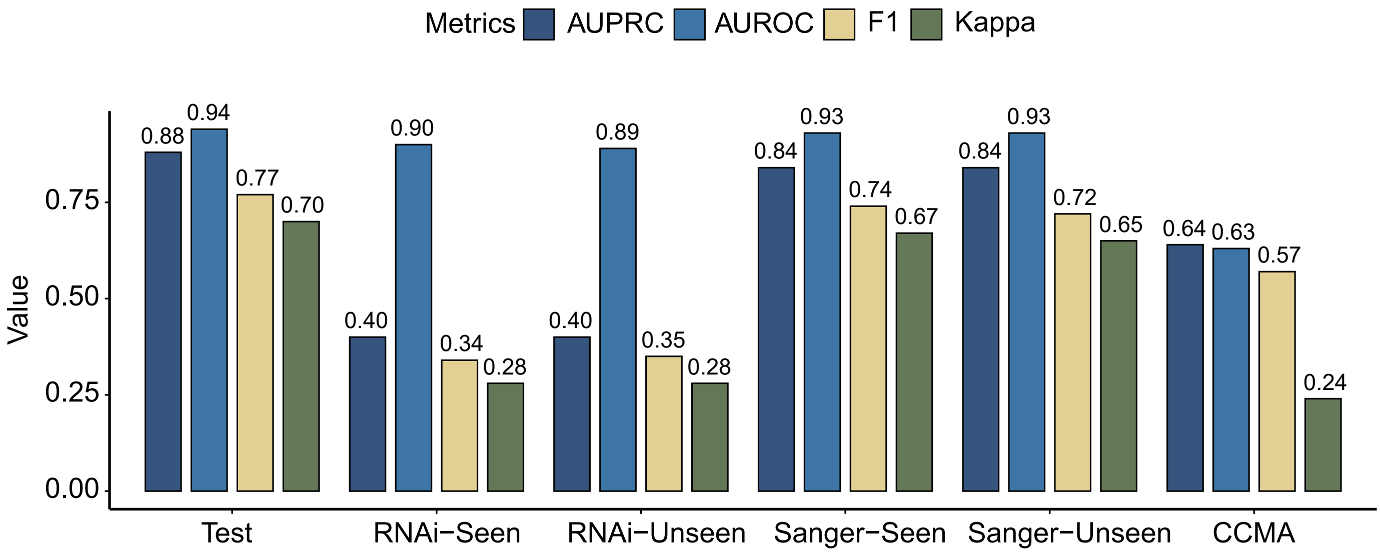


**Figure S3. DeepMeta validation using independent datasets.** DeepMeta performance shown as different metrics on the test set and additional validation datasets, including RNAi dataset, Sanger dataset and CCMA dataset. In RNAi and Sanger dataset, performance in both overlapped and unique cell lines with training dataset were shown.

A

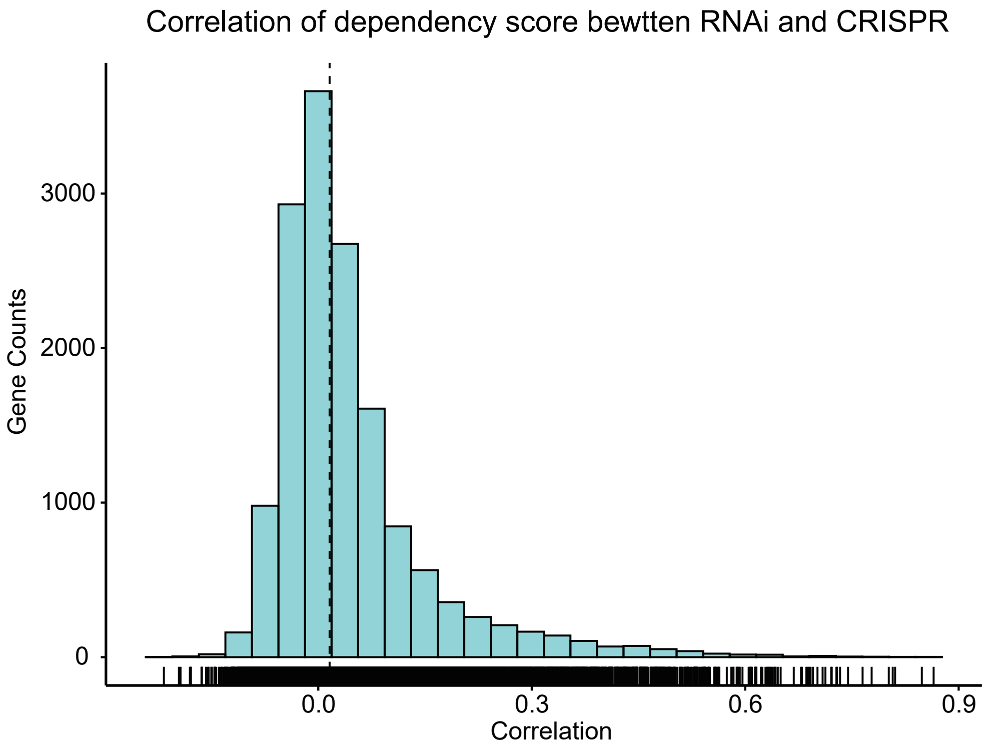


B

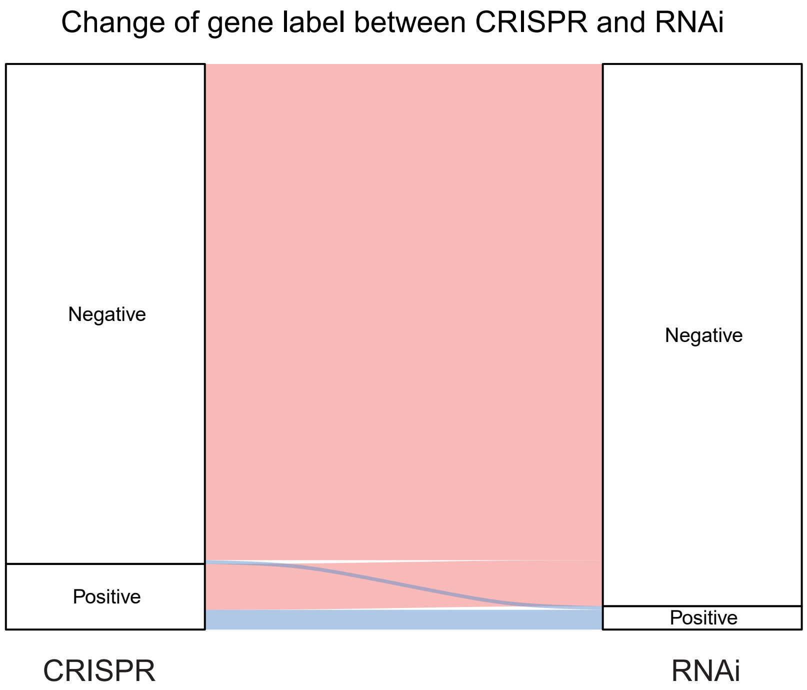


**Figure S4. Comparison between RNAi and CRISPR datasets.** A, The correlation distribution of dependency scores for metabolic gene-cell pairs shared between RNAi and CRISPR datasets. B, Comparing the labels of metabolic genes between the two data sets, the left bar represents the CRISPR data set and the right bar represents the RNAi data set. It can be seen that most of the genes with positive labels in CRISPR dataset are negative in the RNAi dataset.

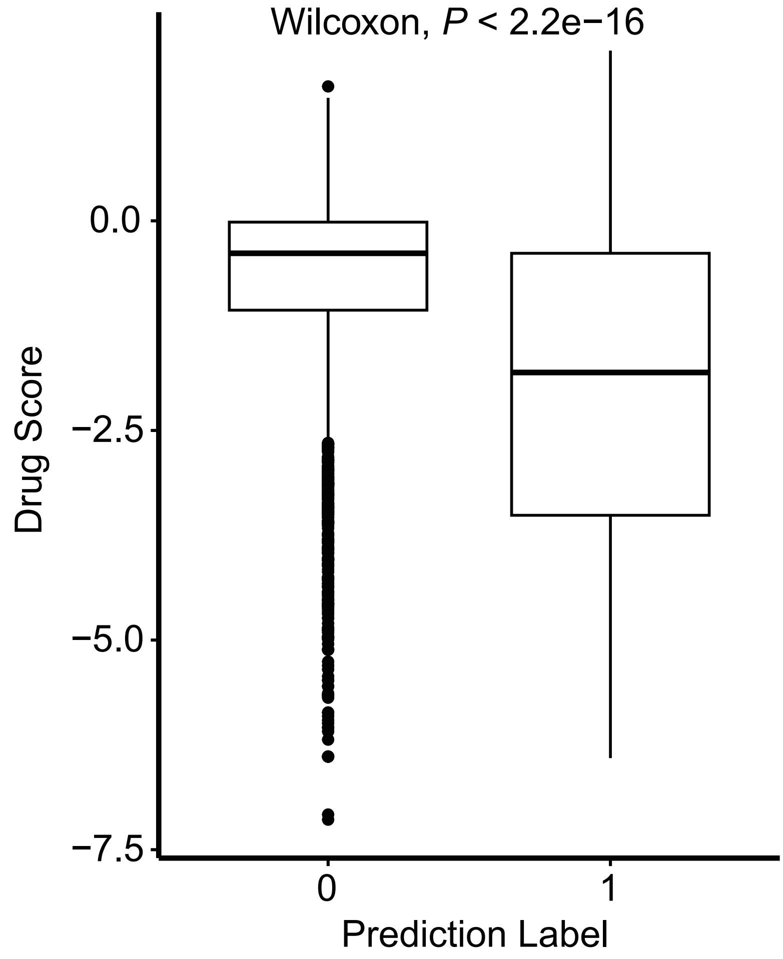


**Figure S5. Validation in PRISM repurposing dataset.** We downloaded primary PRISM Repurposing dataset from DepMap. The drug effect score is represented by log fold change (cell viability between treatment and control).We compared the effect score between drugs targeting genes with predicted positive dependency label and negative dependency label in test dataset (48 cell lines).

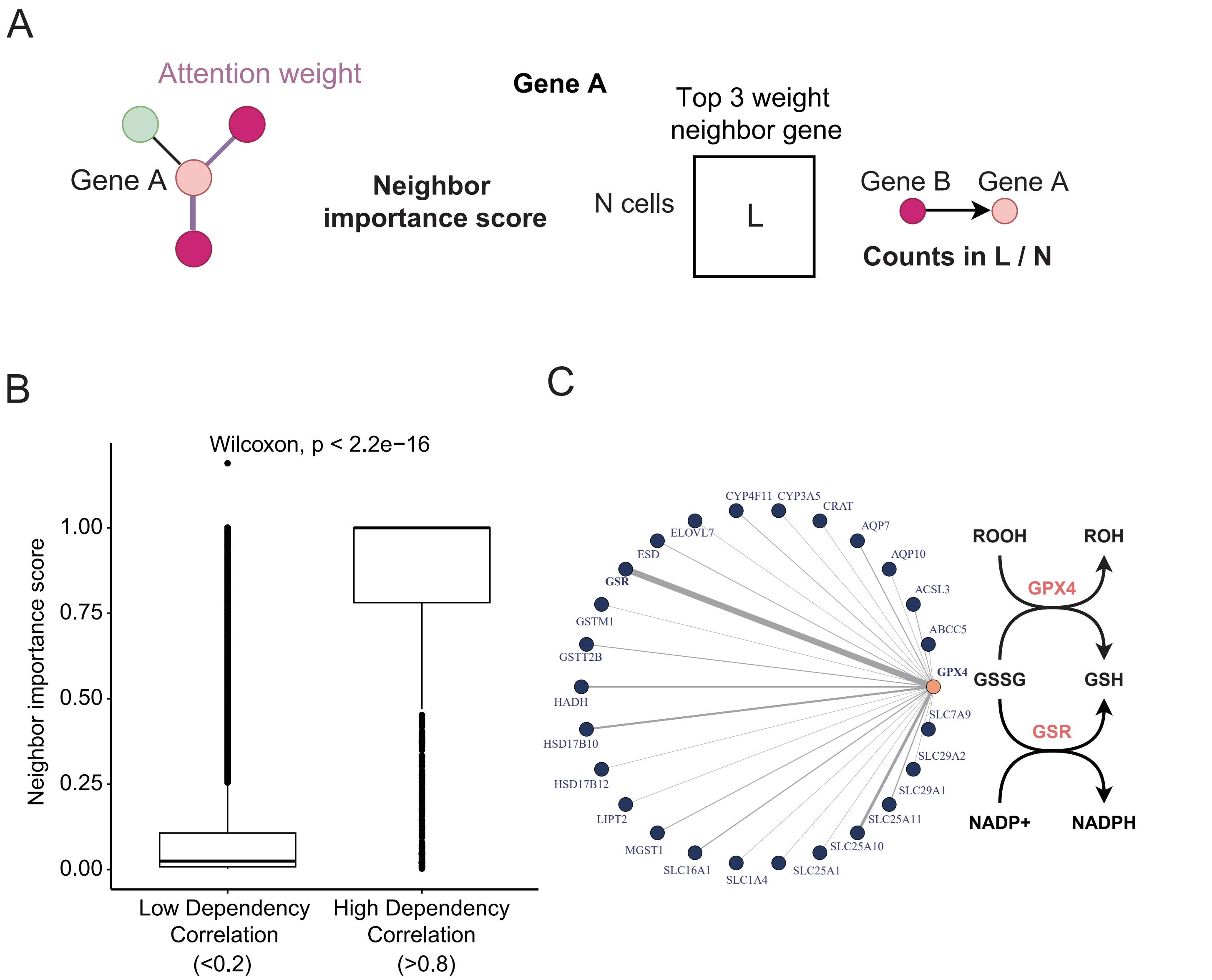


**Figure S6. DeepMeta model interpretation.** A, Left: workflow of model interpretation by neighbor attention. Attention weights from the GAT model were extracted to determine the importance of a gene's neighbor genes for its prediction. Right: definition of the neighbor importance score. For one specific gene A, by extracting the top three attention-weighted neighbor nodes of the gene in the enzyme network of each cell, a matrix of N * 3 is obtained.Neighbor importance of gene B to gene A was calculated as the frequency of gene B appearing in this N*3 matrix divided by the total number of cell lines (N). B, Comparison of neighbor importance scores between high (R > 0.8) and low (R < 0.2) dependency correlation gene pairs. C, Contribution of neighbor genes for predicting the metabolic dependency of the GPX4. The thickness of the edge indicates the neighbor importance score.

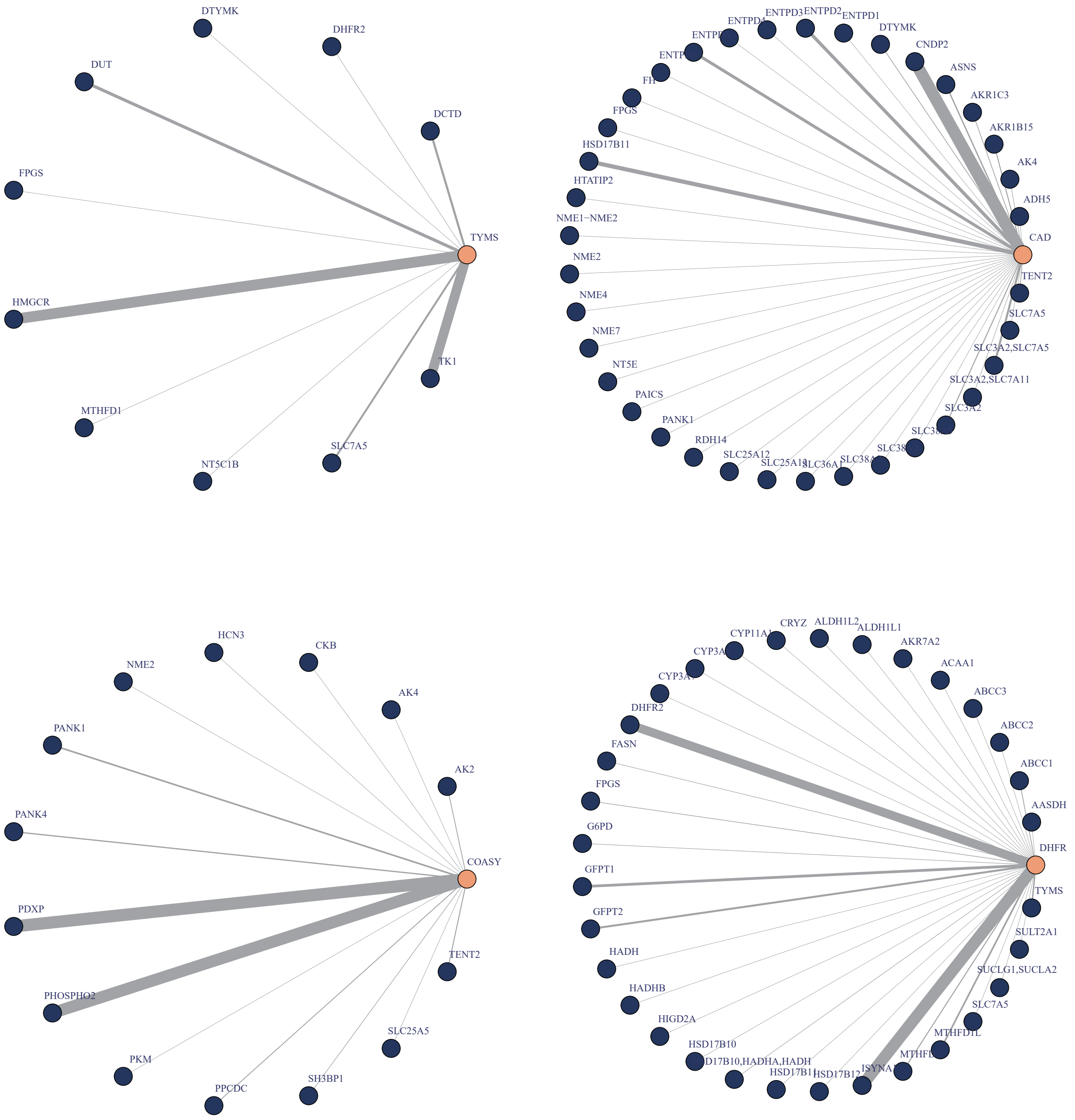


**Figure S7. Examples of neighbor importance interpretation for four metabolic genes TYMS, CAD, COASY and DHFR. The thickness of the edge indicates the neighbor importance scores.**

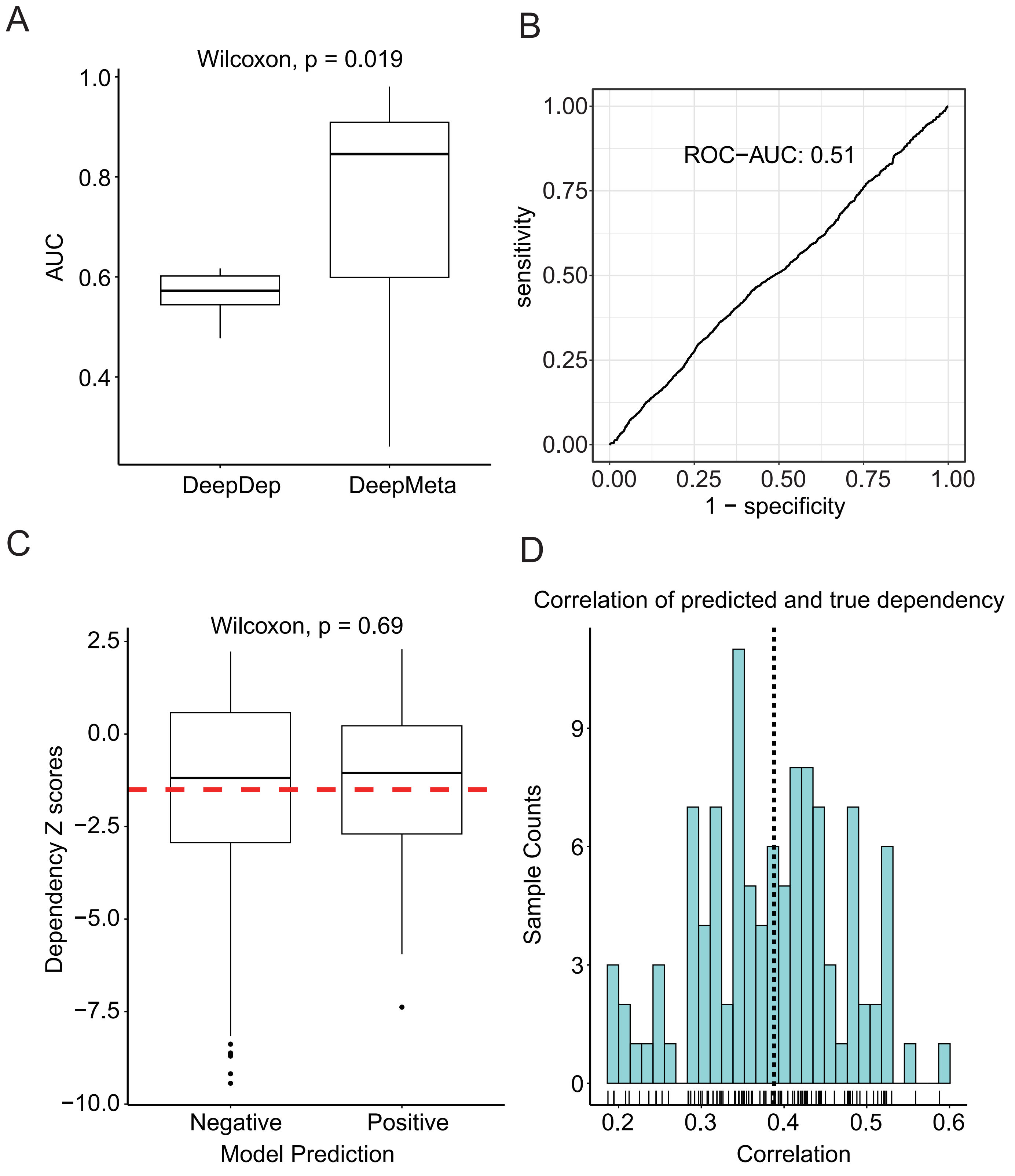


**Figure S8. Performance of DeepDep in drug screen and CCMA dataset.** A,Comparison of model performance between DeepDep and DeepMeta in Pemovska drug screen dataset. In each cell, the gene was assigned a rank value based on AUC score of drugs targeting this gene. We extracted the overlap genes between top 5 genes predicted by models (DeepDep or DeepMeta) and the genes with top 3 AUC rank in each cell. Then the AUC scores of these overlap genes were compared between DeepDep and DeepMeta. B, DeepDep performance in CCMA dataset is shown, and the performance of DeepMeta is shown in Figure 2d left. C, Comparison of Z scores between different categories predicted by DeepDep. The value DeepDep predicted is not probability value, but gene effect scores. Therefore,according to the instructions on the DepMap official website, samples with predicted values less than -0.5 are labeled as positive, and others as negative.The performance of DeepMeta is shown in Figure 2d right. D, Distribution of correlation of predicted dependency and z scores.

A

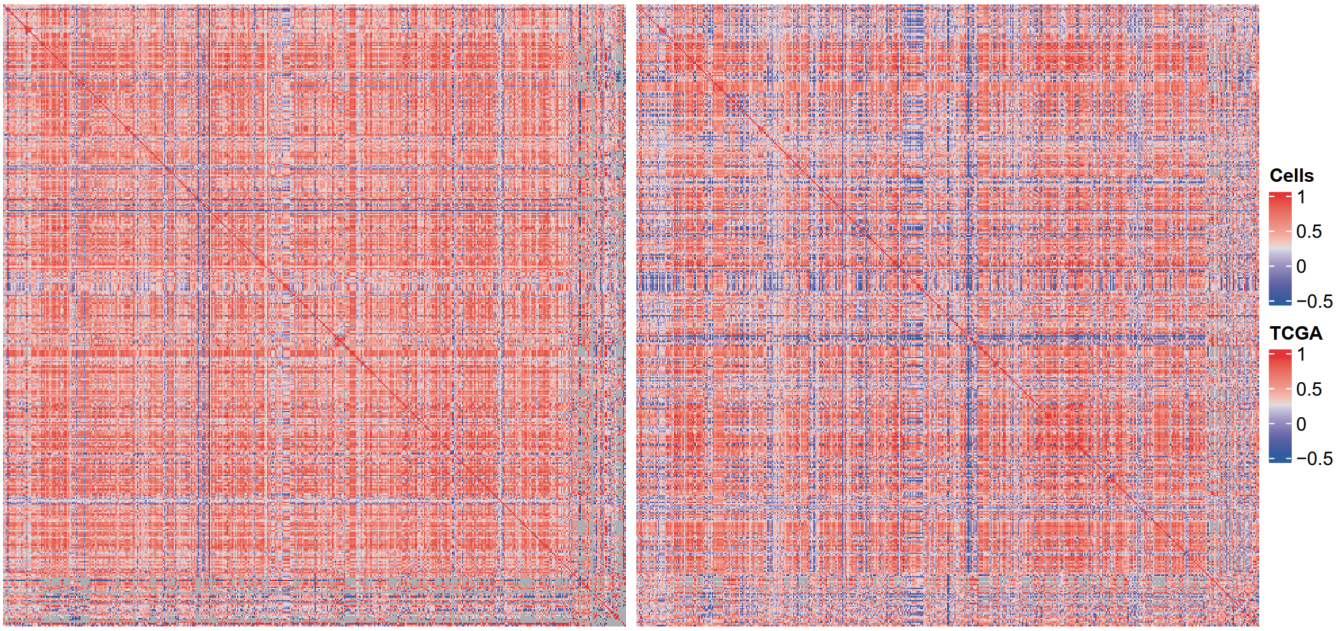


B

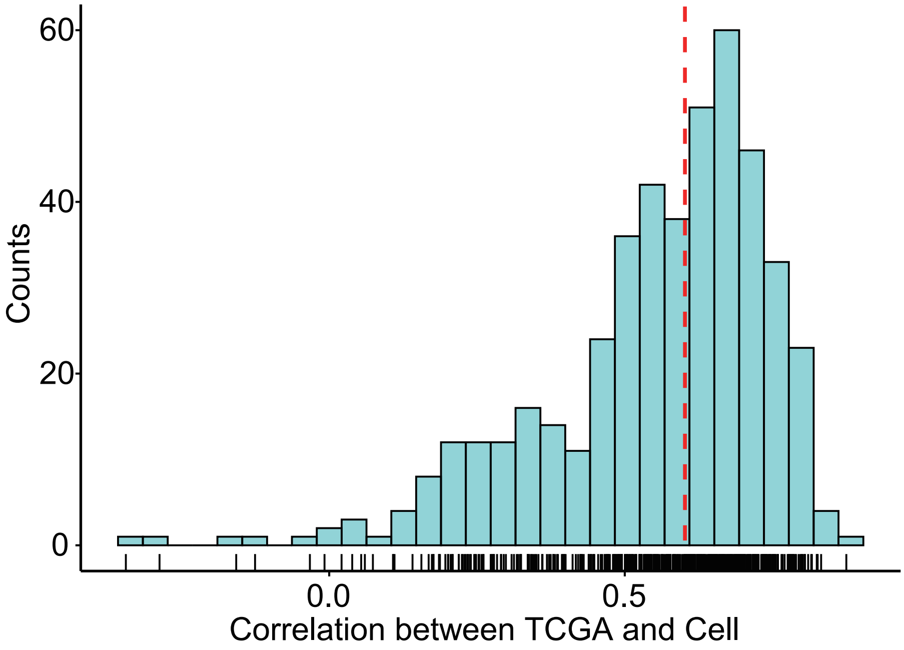


**Figure S9. Consistency of gene dependency predictions in TCGA tumor samples and DepMap cell lines.** A, Spearman correlation between prediction dependency of pairwise genes in the cell line data (left) and TCGA (right) data respectively. B, Spearman correlation between the corresponding columns of the cell line matrix and the TCGA tumor sample matrix in (A).

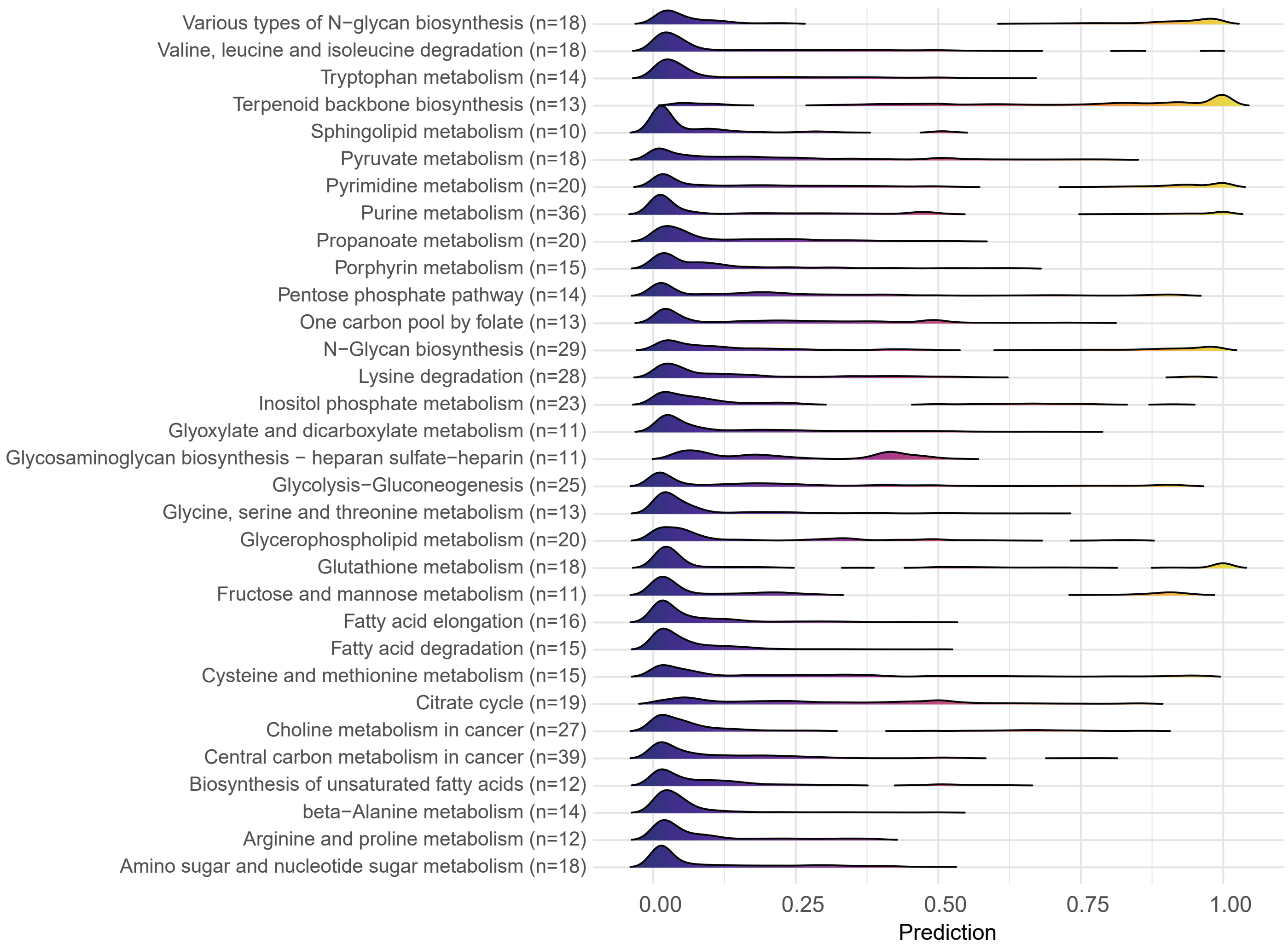


**Figure S10. Distribution of predicted metabolic dependency scores of genes in KEGG metabolic pathways by applying DeepMeta in TCGA samples, only pathways with more than 10 genes are shown.**

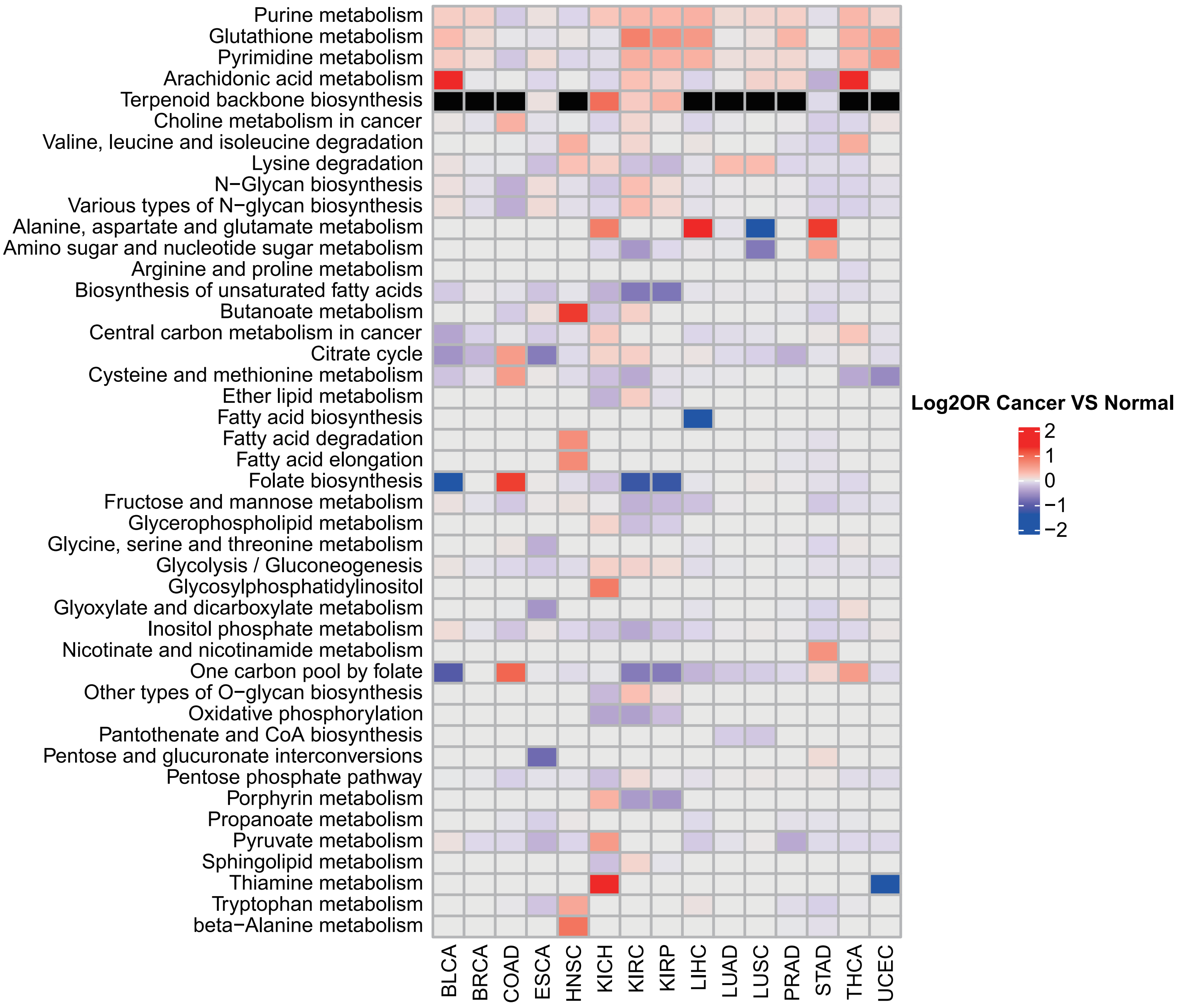


**Figure S11. Comparison of metabolic pathway dependency odds ratio (OR)values (obtained by one tail Fisher test) between TCGA cancer and corresponding normal samples.** The value in the heatmap is the median log2OR of the pathway in cancer tissue minus the median log2OR in normal tissue. Only TCGA cancer types with more than 10normal samples are shown.

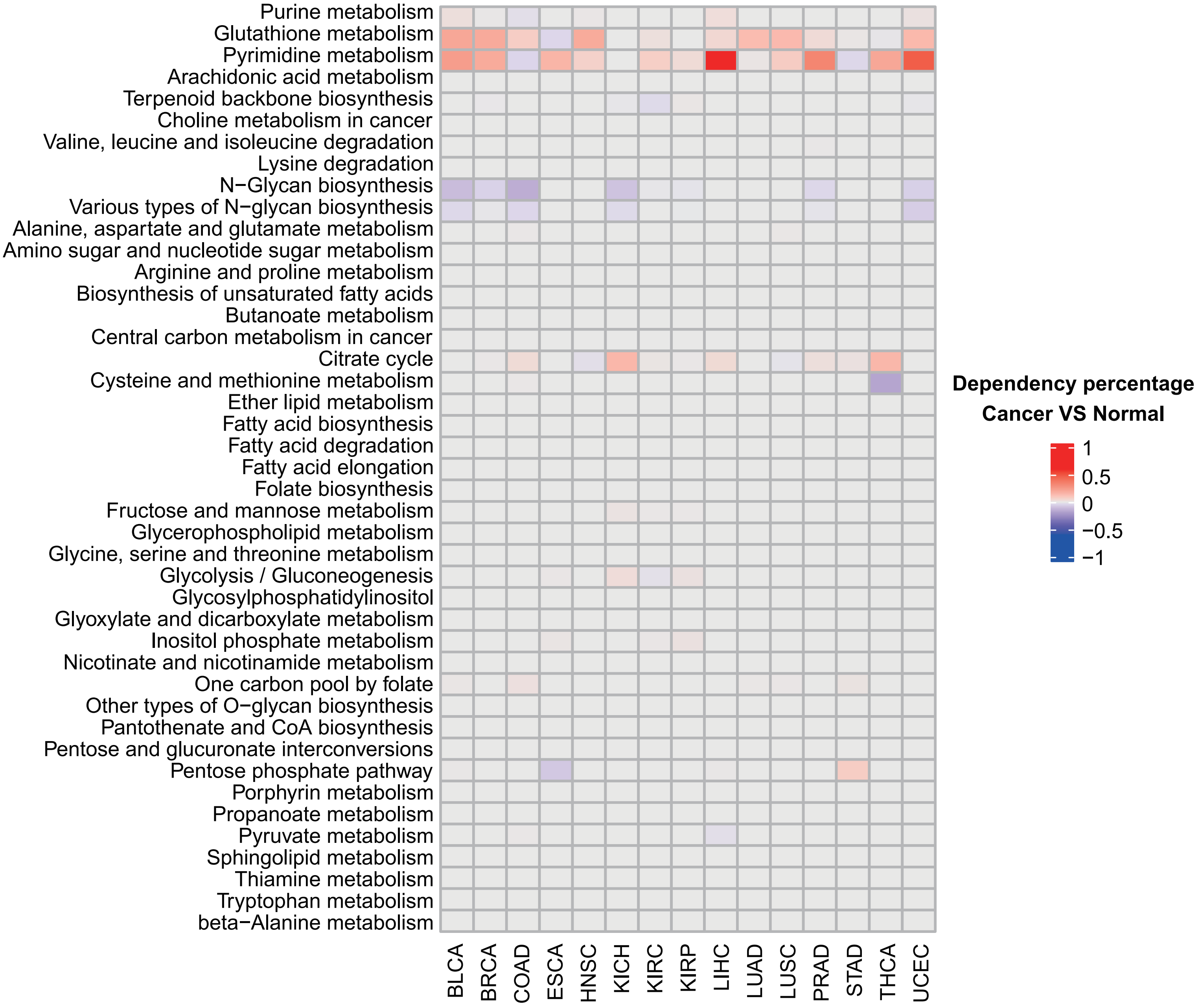


**Figure S12. Comparison of metabolic pathway dependency between TCGA cancer and corresponding normal samples.** Values indicate difference of percentage of samples show dependency to the indicated metabolic pathway between TCGA cancer and corresponding normal tissues. The dependency percentage was determined by permutation test. For a given pathway in a particular sample, we randomly sampled genes equal to the number of genes in that pathway to form a "random gene set". We calculated the proportion of genes predicted as positive in this random gene set, repeated this process 1000 times to obtain 1000 proportion values. Then, we compared this empirical distribution of 1000 values with the actual proportion of positive genes in the pathway to obtain a P-value. Thus, for each sample, we can get such a permutation test P value. If this P value was less than 0.05, then the sample was defined as dependent on this metabolic pathway. For each metabolic pathway, we calculated the proportion of samples with the P-value less than 0.05 in both the cancer tissue and its corresponding normal tissue.Subsequently, we determined the difference by subtracting the proportion in the cancer from the proportion in the normal tissue, which yielded the values displayed in the heatmap.

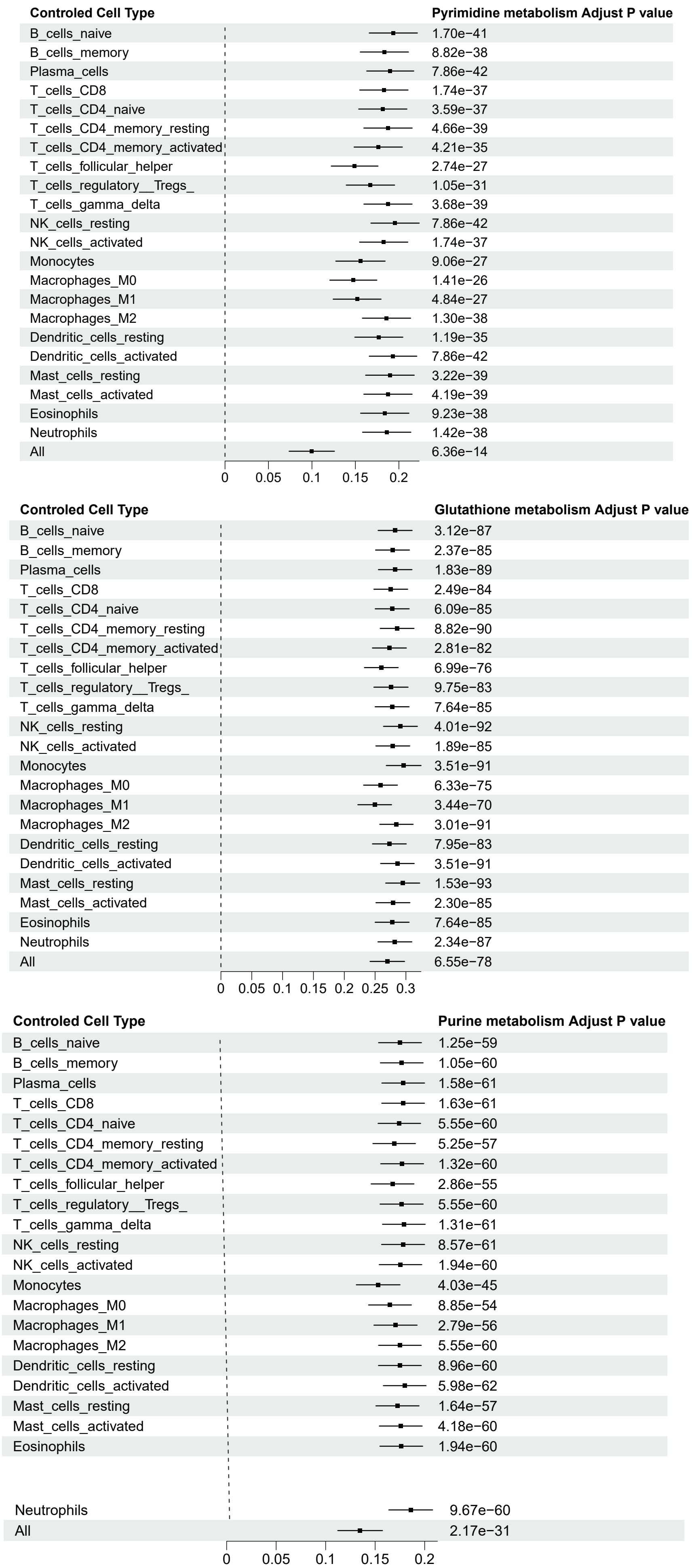


**Figure S13. Compare dependency of three metabolic pathway in cancer vs normal samples in TCGA pan-cancer dataset when control for different stromal cell abundance.** We utilized linear regression to model the dependency of metabolic pathway and sample type (cancer or normal),including different stromal cell abundance as covariates: Dependency ~Sample-Type + Cell-Abundance. The dependency of the pathway was quantified by log odds ratio value same as Figure3A, and the stromal cell abundance was quantified by CIBERSORT. The coefficients, adjust p value and corresponding 95% confidence level of sample type variable were shown in forest plot

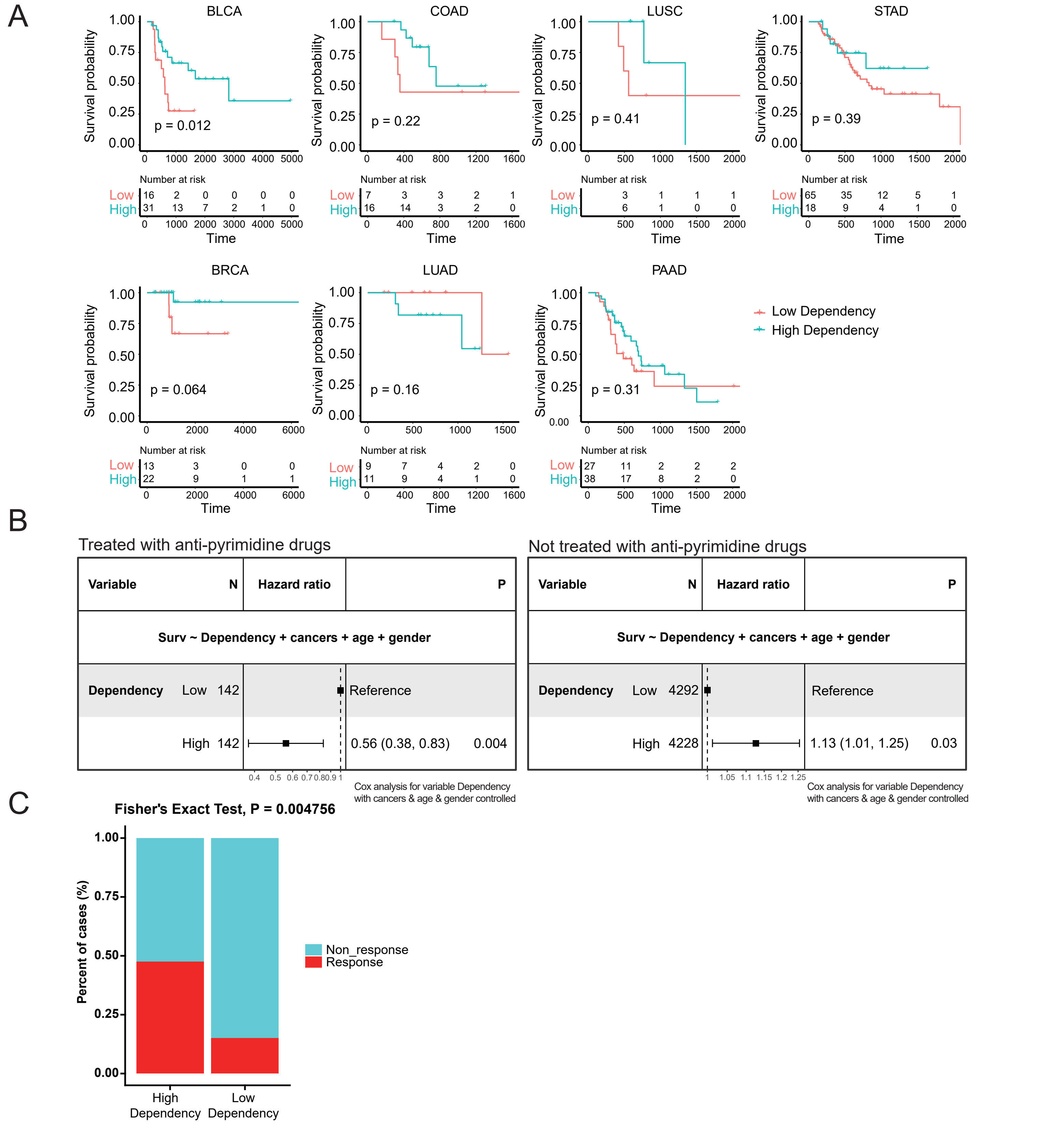


**Figure S14. Survival analysis of TCGA samples with different pyrimidine metabolism dependency treated or not treated with anti-pyrimidine drugs.** A, Kaplan-Meier overall survival curves show the comparison between different patient groups stratified by pyrimidine metabolism pathway dependency status in different cancer types. The dependency of pyrimidine metabolism pathway was calculated by one tail Fisher test and higher odds ratio (OR) value indicates that samples are more dependent on pyrimidine metabolism pathway, compared to all other pathways (Methods). Only cancer types with sample counts more than 10 were shown. B, Cox proportional hazard analysis considering cancer type, age and gender as covariate is shown for TCGA samples treated with four anti-pyrimidine drugs including capecitabine, pemetrexed, gemcitabine and fluorouracil (left panel), or in all TCGA samples except those treated with anti-pyrimidine drugs (right panel).C, Validation of DeepMeta clinical efficacy in anti-pyrimidine drug chemotherapy cohort. We downloaded gene expression and treatment response data from CTR-DB database. These patients were treated by fluorouracil based anti-pyrimidine drugs and outcome was recorded as response or non-response. DeepMeta was used to predict dependency of pyrimidine metabolism pathway. The cutoff between “high” and “low”dependency patient group is the top 20% and bottom 20% dependency score for pyrimidine metabolism pathway dependency.

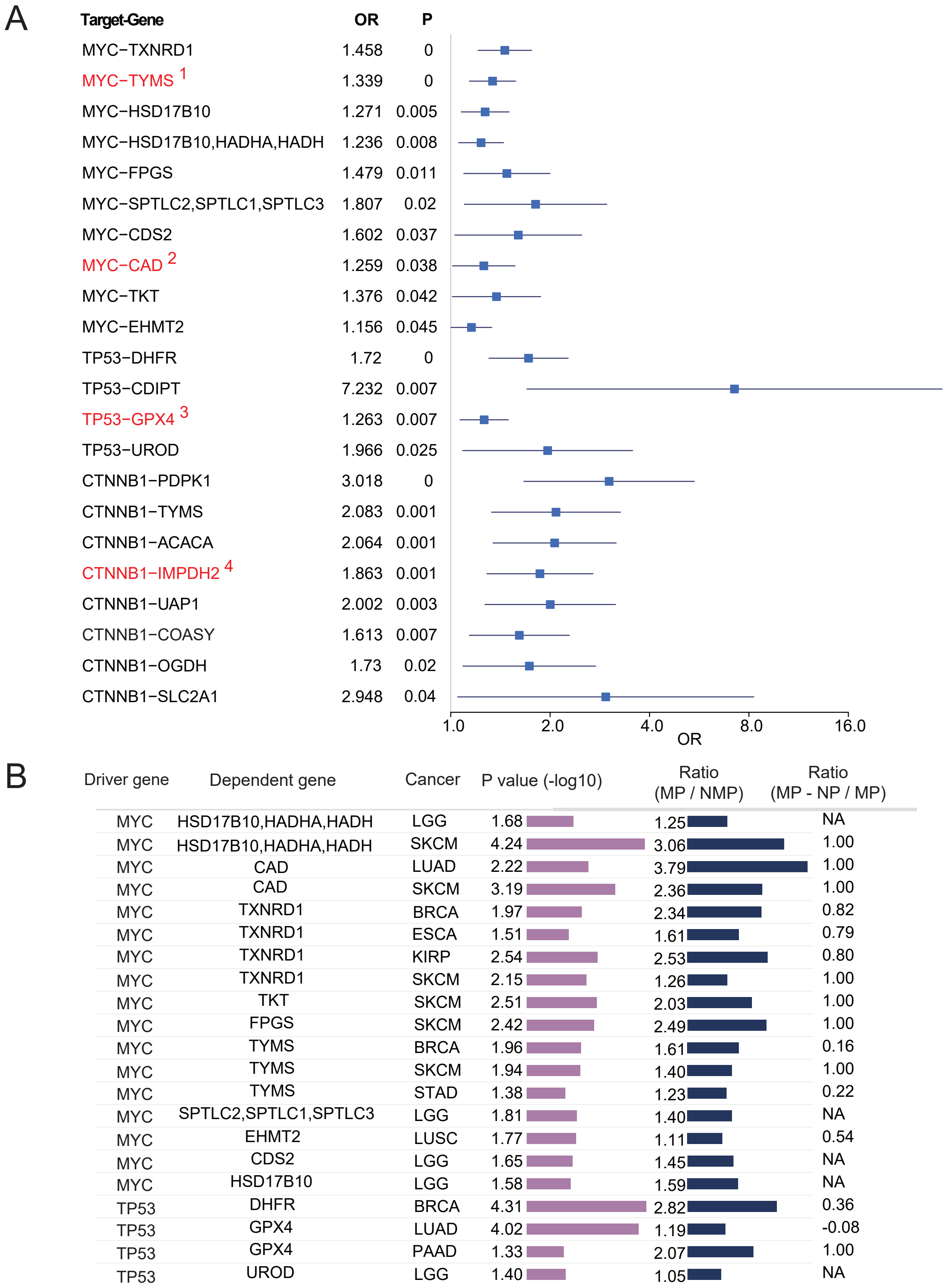


**Figure S15. Metabolic dependencies prediction for undruggable cancer driving genetic alterations.** A, Pan-cancer metabolic target screen results of three cancer driver genes. The odds ratio (OR) and P values were calculated by generalized linear model which models the relationship between predicted metabolic dependencies and mutation status of cancer driver genes and controls for cancer type variable. For a specific metabolic gene, the OR value of the mutation status variable in the generalized linear model is the ratio of the proportion of samples predicted to be dependent on this metabolic gene in the mutant samples and proportion of samples predicted to be dependent on this metabolic gene in the non-mutated samples. Thus, only genes with OR greater than 1 and P value < 0.05 are shown here. The metabolic targets of some cancer driving genes have been reported in literature and marked in red,and the related references have been indicated: 1. Liu. et.al1, 2. Miltenberger.et.al19, 3. M Tahaney. et.al4, 4. Chong. et,al20. B, Summary of the identified metabolic dependencies of cancer driver genes. P values are calculated by one tail Fisher test. MP: Proportion of predicted positive cases in cancer samples with functional mutations in the driver gene, NMP: Proportion of predicted positive cases in cancer samples without mutations in the driver gene, NP: Proportion of predicted positive cases in normal samples.

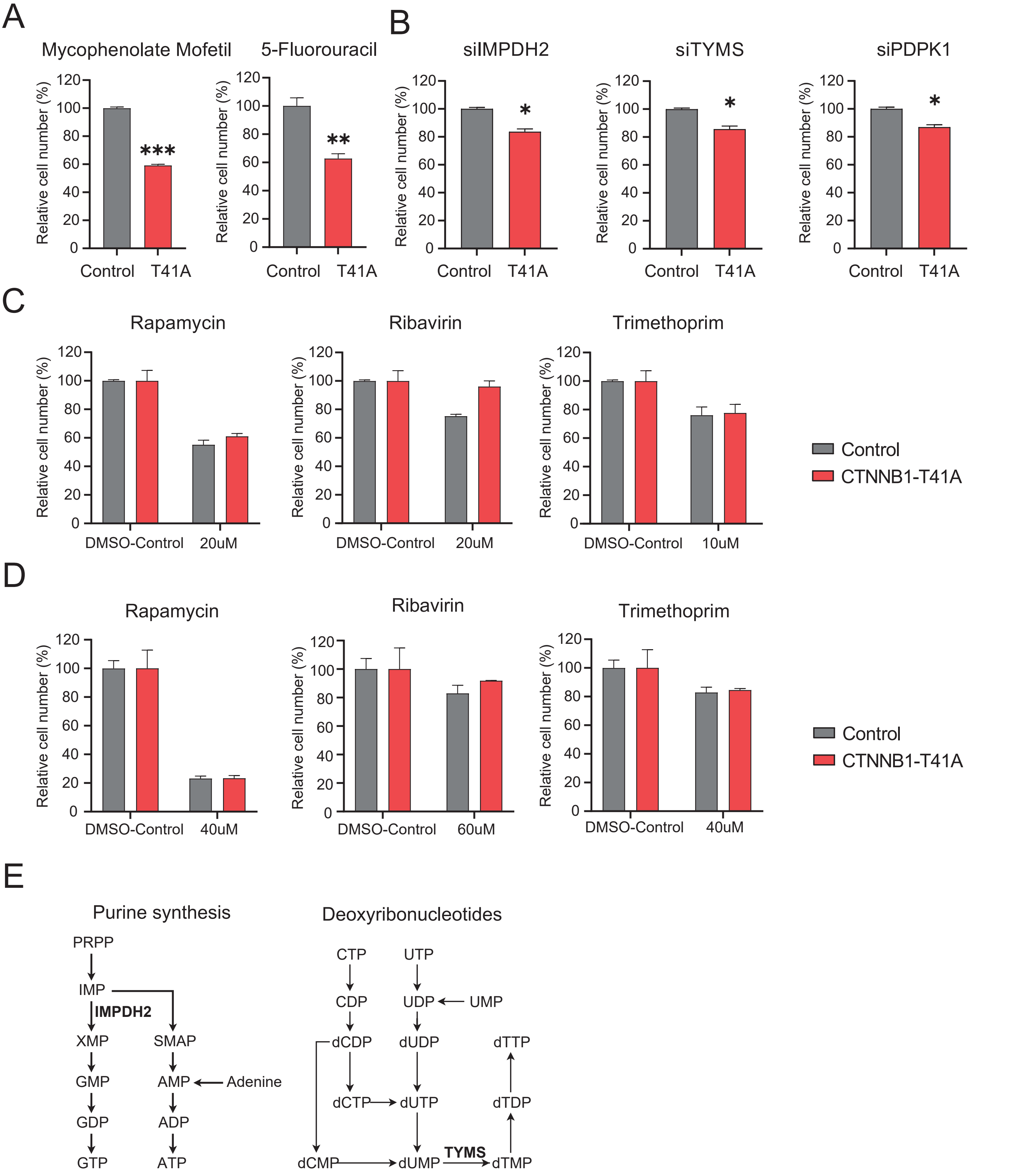


**Figure S16. Experimental validation of predicted metabolic dependency targets in CTNNB1-mutant cell lines.** A, Effects of mycophenolate mofetil (IMPDH2 inhibitor, 5uM) and 5-fluorouracil (TYMS inhibitor, 20uM) on cell growth in CTNNB1-T41A mutant 293T cell lines. B, Effects of shRNA-mediated knockdown of PDPK1, TYMS, and IMPDH2 in the growth of CTNNB1-T41A mutant 293T cells. C-D, Non-selective effects of random drugs on the growth of CTNNB1-T41A WI38 (C) and 293T (D) cell lines. Control:cells infected with a lentivirus carrying the vehicle control; CTNNB1-T41A:cells stably expressing the CTNNB1-T41A mutation. E, Schematic representation of the nucleotide metabolism pathway involving IMPDH2 and TYMS.

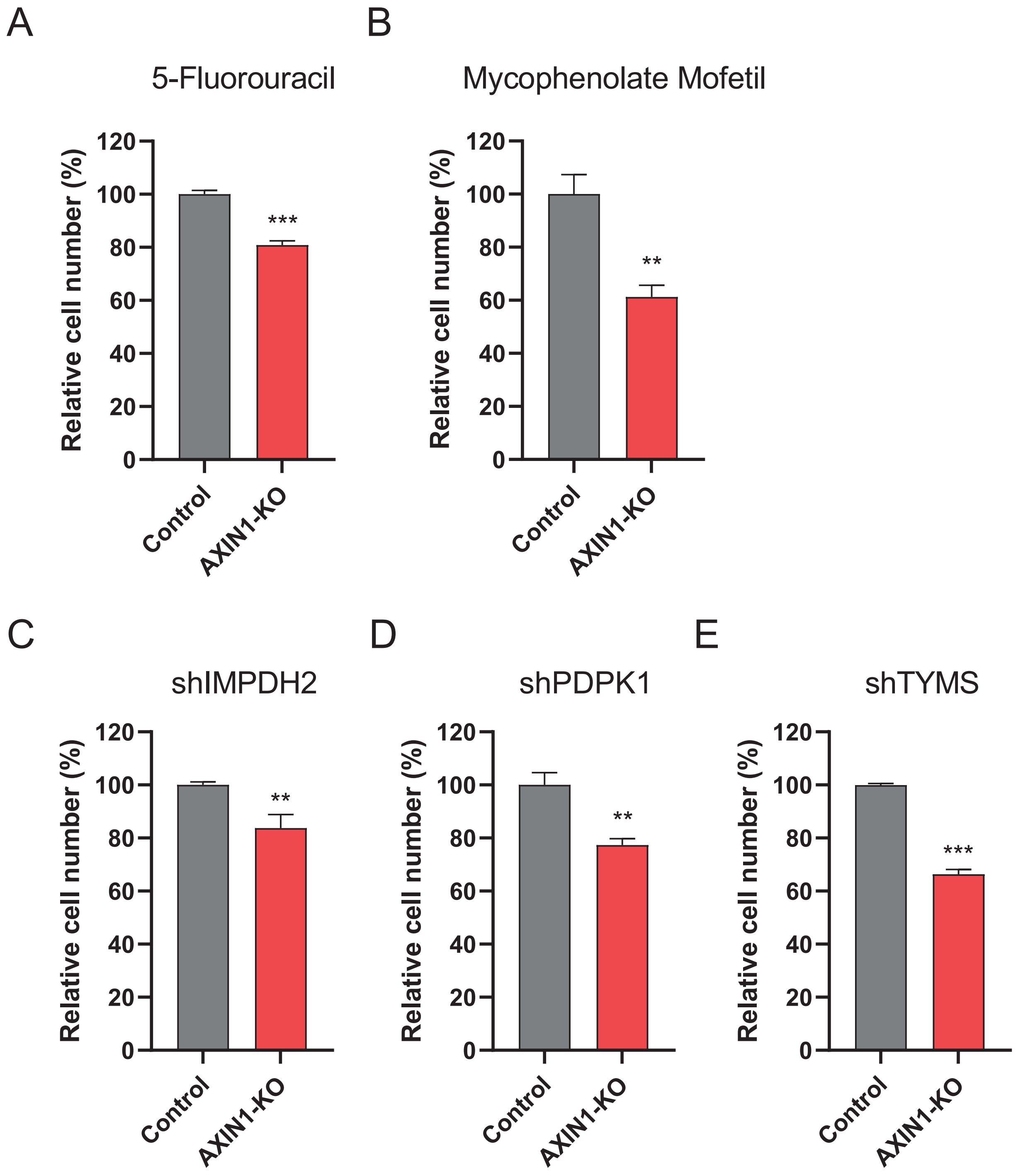


**Figure S17. Experimental validation of predicted metabolic dependency targets in AXIN1 knockout cell lines.** The viability (%) of AXIN1-knockout and control MHCC97-H cells treated with 5 μM 5-Fluorouracil (A) and 60 μM mycophenolate mofetil (B), respectively, was measured. Additionally, cell proliferation rate (%) was assessed for control and AXIN1-knockout MHCC97-H cells with shRNA-mediated knockdown of IMPDH2 (C), PDPK1 (D),and TYMS (E). The control cells were sgRNA control cells.

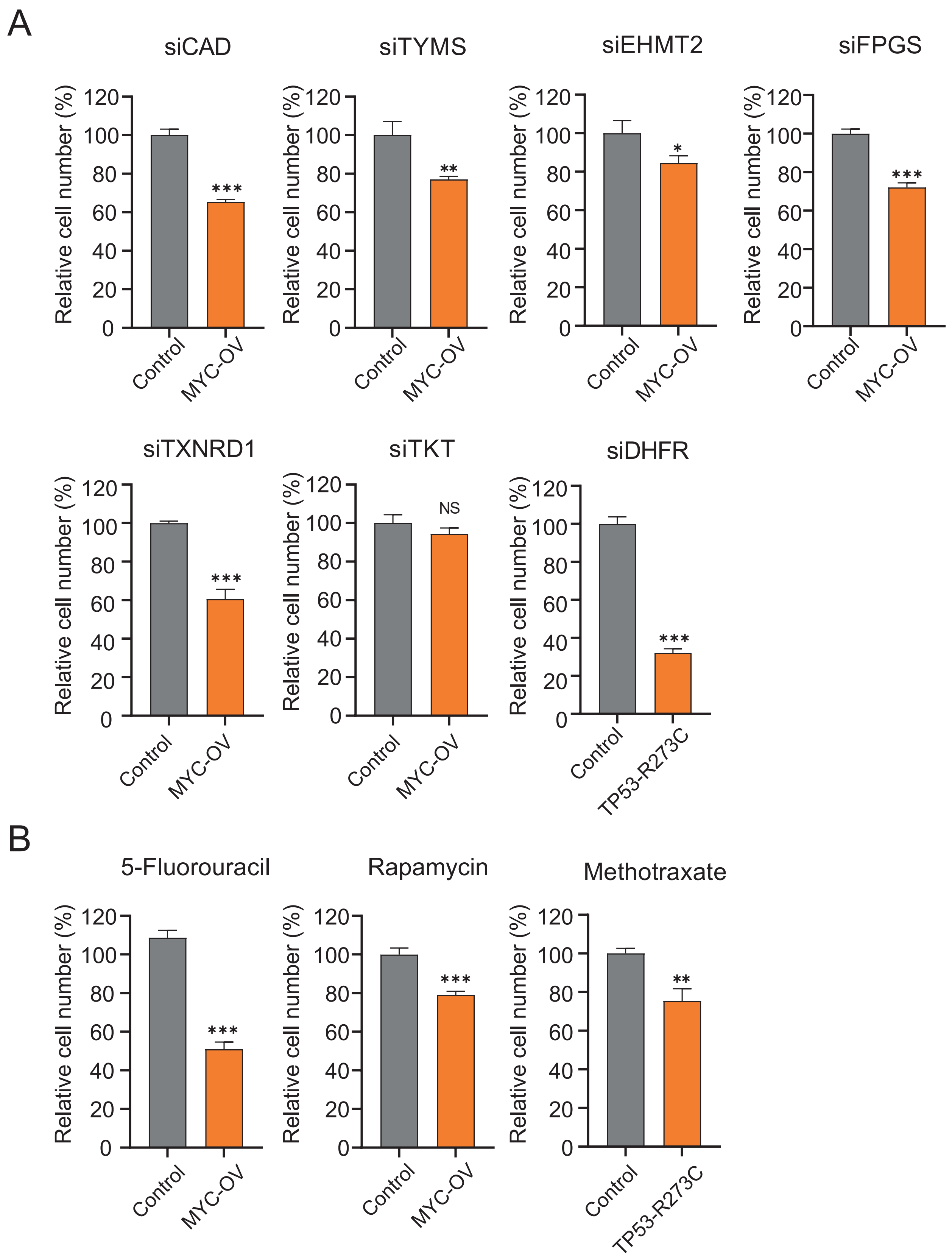


**Figure S18. Experimental validation of predicted metabolic dependency targets in cell lines with MYC overexpression, TP53 inactivating mutation.** siRNA-mediated gene knockdown (A), and chemicals-mediated target inhibition (B) validated the metabolic dependency targets predicted by DeepMeta in cell lines with overexpressed MYC (MYC-OV), or TP53 inactivating mutation (TP53-R273C). Control: cell lines carrying the vehicle.

### Table S1. siRNA sequence information targeting genes of interest

| Genes | siRNA-sense | siRNA-antisense |
|---|---|---|
| siNC | UUCUCCGAACGUGUCACGUTT | ACGUGACACGUUCGGAGAATT |
| PIGC | CCCUGCAUGCCUUCAUCAU | AUGAUGAAGGCAUGCAGGG |
| CAD | GCUAGCUGAGAAAAACUUU | AAAGUUUUUCUCAGCUAGC |
| EHMT2 | GCAAAUAUUUCACCUGCCA | UGGCAGGUGAAAUAUUUGC |
| FPGS | AGAGCUAAAGAAUCCCGUC | GACGGGAUUCUUUAGCUCU |
| TKT | CCAGCCAACAGCCAUCAUUTT | AAUGAUGGCUGUUGGCUGGTT |
| TXNRD1 | AAUGAUAGAAGCUGUACAGAA | UUCUGUACAGCUUCUAUCAUU |
| DHFR | GUCUAGAUGAUGCCUUAAA | UUUAAGGCAUCAUCUAGAC |

### Table S2. Drug information targeting genes of interest

| Genes | Drugs | Brand | Catalog number |
|---|---|---|---|
| IMPDH2 | Mycophenolate Mofetil | Solarbio | IM0820 |
| TYMS | 5-Fluorouracil | Cayman chemical | 14416 |
| CAD | Rapamycin | Adamas-beta | 62975G |
| DHFR | Methotrexate | Solarbio | IM0160 |
| | Trimethoprim | Selleck | S3129 |
| AHCY | Ribavirin | Solarbio | IR0090 |
